# Picomolar-Level Sensing of Cannabidiol by Metal Nanoparticles Functionalized with Chemically Induced Dimerization Binders

**DOI:** 10.1101/2023.09.13.557660

**Authors:** MD Ashif Ikbal, Shoukai Kang, Xiahui Chen, Liangcai Gu, Chao Wang

## Abstract

Simple and fast detection of small molecules is critical to health and environmental monitoring. Methods for chemical detection often use mass spectrometers or enzymes; the former relies on expensive equipment and the latter is limited to those that can act as enzyme substrates. Affinity reagents like antibodies can target a variety of small-molecule analytes, but the detection requires successful design of chemically conjugated targets or analogs for competitive binding assays. Here, we developed a generalizable method for highly sensitive and specific in-solution detection of small molecules, using cannabidiol (CBD) as an example. Our sensing platform uses gold nanoparticles (AuNPs) functionalized with a pair of chemically induced dimerization (CID) nanobody binders (nano-binders), where CID triggers AuNPs aggregation and sedimentation in the presence of CBD. Despite moderate binding affinities of the two nano-binders to CBD (*K*_D_s of ∼6 and ∼56 µM), a scheme consisting of CBD-AuNP pre-analytical incubation, centrifugation, and electronic detection (ICED) was devised to demonstrate a high sensitivity (limit of detection of ∼100 picomolar) in urine and saliva, a relatively short assay time (∼2 hours), a large dynamic range (5 logs), and a sufficiently high specificity to differentiate CBD from its analog, tetrahydrocannabinol. The high sensing performance was achieved with the multivalency of AuNP sensing, the ICED scheme that increases analyte concentrations in a small assay volume, and a portable electronic detector. This sensing system is readily coupled to other binders for wide molecular diagnostic applications.

## 1. Introduction

Small molecules are ubiquitous, taking the form of metabolites, drugs, toxins, etc.^1–3^, and have been explored for pain relief ^4^ and treatment of cardiac diseases^5^, cancer^6^, and infectious diseases^7^ including COVID-19 ^8^. It is critical to differentiate relatively subtle differences in small molecules, because they can encipher drastic differences in the molecular bioactivities. For example, delta-9-tetrahydrocannabinol (THC) and cannabidiol (CBD)^9, 10^ are two major constituents of cannabis with similar structures but different pharmacology and psychoactivity. The current gold standard detection method is mass spectrometry^11^, which, however, is commonly used in centralized facilities instead of standardly equipped laboratories. Portable methods such as electrochemical glucose sensors ^12^ are highly useful, but the detection relies on the availability of a specific enzyme that recognizes the small molecule substrate and thus is not widely applicable to any analyte. In comparison, affinity-based detection methods, such as an enzyme-linked immunosorbent assay (ELISA)^13, 14^ are complementary and more generalizable, but they typically require the labeling of small molecules for competitive binding assays, i.e. chemical conjugation of small molecule targets or their competitive analog to solid support or reporter molecules^15^. In this regard, chemically induced dimerization (CID)^16, 17^ provides an ideal mechanism to directly sense small molecules via forming a ternary complex, making it suitable for in-solution detection without washing steps. Recently, we developed a method (COMBINES-CID) for creating CID systems by screening a combinatorial nanobody library applicable to different analytes^18^. CID binders for CBD were selected^18^ and found to dimerize via a “molecular glue” mechanism^19^. Although ELISA-like sandwich assay was applicable for CBD detection, such assay required extensive sample incubation, washing, and spectrometric or fluorescent analysis, thus not suitable for portable detection. It is therefore necessary to develop a simple, fast sensing platform with a digital readout for a variety of analytes.

Here, we report a sensitive and rapid detecting system with an inexpensive optoelectronic readout that significantly improves the detection limit (by eight times) and reduces the detection time (from ∼5 hours to <2 hours) compared to ELISA with the same CID nano-binder pair. Specifically, multivalent sensors were designed by conjugating synthetic nanobinders specific to the small molecules onto gold nanoparticles (AuNPs). Such nano-binders can dimerize with the small molecules, thus triggering aggregation and subsequent precipitation of AuNP sensors. This accordingly results in increased solution transparency correlated with small molecule concentration. The modulation of solution color can be further quantified using a lab-based spectrometer or a portable optoelectronic readout system comprising simple and inexpensive components such as light-emitting diode (LED), photodiode, and a battery. Using CBD as the target, we demonstrate a new scheme consisting of CBD incubation with binder-functionalized AuNPs, <u>c</u>entrifugation and <u>e</u>lectronic <u>d</u>etection (ICED) to achieve a high sensitivity (<100 pM) in urine and saliva (roughly 3 logs lower than the typical concentration in human body^16^), a high specificity (distinguishing from CBD homologue THC) and a large dynamic range (5 logs). This ICED scheme strongly facilitates high-performance small molecule recognition by the multivalent functionalization of binder pairs on AuNPs, localized detection of concentrated CBD via incubation on AuNPs and centrifugation, accelerated signal transduction from AuNP aggregation and precipitation, and low background noise from portable electronic detectors. This cost-effective, portable and accurate system can be advantageous in broad applications such as drug and toxin detection, biomarker diagnostics, drug discovery, etc.

## 2. Results and Discussion

### 2.1 CBD Binder Protein Selection

To obtain specific binders to the CBD molecules, a COMBINES-CID method wa employed^18, 20^ to design the CBD CID system, isolated from a combinatorial nano-binder library of over 10^9^ complementarity-determining region (CDR) peptide sequences. To obtain CBD anchor binder (CA), six rounds of selection were performed using biotinylated CBD as bait, eventually obtaining three unique clones with high specificity. CA14, a binder with the highest protein yield was then chosen as bait for dimerization binder (DB) selection, producing 24 unique DBs after four rounds of biopanning. The two most stable CBD CID binder pairs, CA14-DB21(Kd= 56 nM) and CA14-DB18 (Kd= 560 nM), were expressed as a C-terminal Avi-tagged and His-tagged form in E. coli, purified by Ni-affinity and biotinylated by BirA and then site-specifically conjugated to gold nanoparticles as previously reported^21^.

### 2.2 Preparation of Multivalent Small Molecule Sensor

To prepare for the CBD small molecule assay, we examined AuNPs covalently coated with high-density streptavidin proteins (e.g. estimated ∼1300 binding sites for 80 nm AuNPs) for sensing, conceptually similar to our recently reported antigen sensors^15^. Upon mixing with biotinylated nano-binders (anchor binder CA14 and dimerization binders DB18 or DB21), these AuNPs formed multivalent sensors (Figure 1a). Upon the introduction of small molecules, these multivalent in-solution sensors display improved effective affinity, beneficial for accelerated molecular binding and higher sensitivity compared to monovalent detection systems^22, 23^. The molecular recognition is accompanied by aggregation of the AuNP sensors, resulting in sedimentation. This sedimentation based detection mechanism differs fundamentally from other plasmonic colorimetric assays, where the detection is based on resonance wavelength shift caused by the formation of dimers and oligomers of small AuNP (5 to 20 nm) still present in the solution^24–26^. First, the use of larger AuNPs produces of nanosensors of bigger surface areas, which accordingly hosts more nano-binders and presents a high binding affinity for high-quality multivalent molecular recognition. Further, larger NP sizes also make them more responsive to centrifugation-enhanced target molecule binding and NP precipitation (Figure 1b), which is important to shortening the assay time limit (e.g. demonstrated 2 hours, compared to 5 hours by incubation only) and improving the detection. In addition, our precipitation-enhanced, molecule concentration-dependent modulation of AuNP clustering allows accurate signal readout by probing the optical extinction of free-floating AuNPs. Therefore, the molecular signals can be converted to electronic output by fundamentally a pair of simple LEDs and photodetectors operating at the AuNP extinction wavelengths (Figure 1c), enabling portable, rapid, accurate, inexpensive and digital diagnostics. This method is fundamentally different from and much simpler than conventional optical sensing methods that analyze the complete spectra using bulky and expensive microscopy and spectrometers.

**Figure 1.**
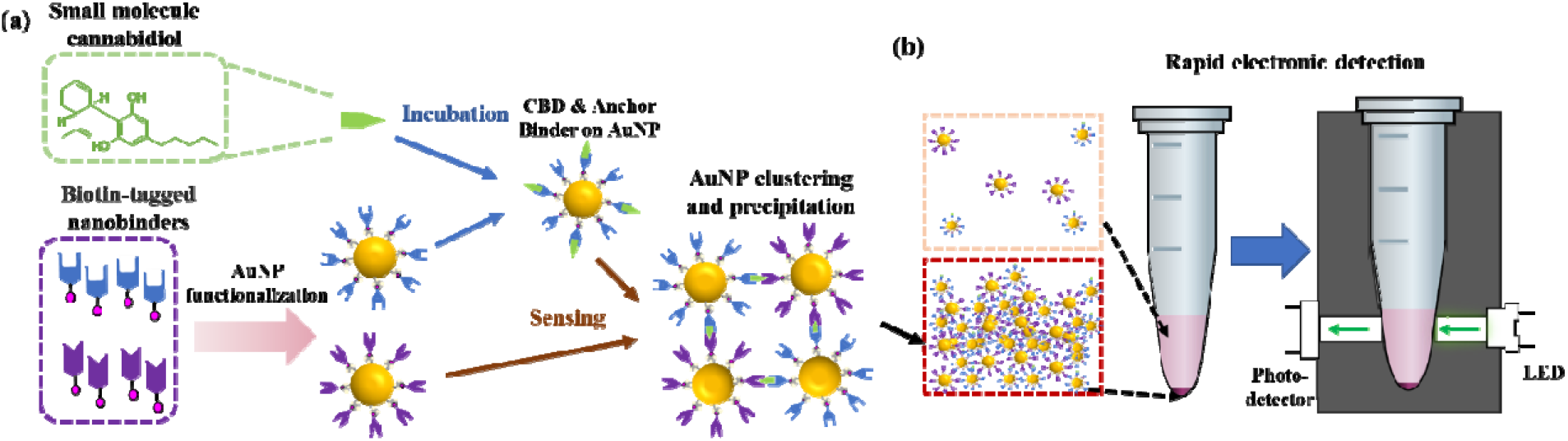
Overview of small molecule detection. (a) Schematic showing key steps in enhanced CBD detection, including functionalization of AuNPs with nano-binders (anchor binder: blue; dimerization binder, purple) (pink arrow), pre-incubation of CBD sample with anchor-binder on AuNPs (blue arrows), and mixing the CBD-anchored AuNPs with dimerization-binder-functionalized AuNPs for reaction (brown arrows). (b) Schematics showing CBD molecule signal readout. Here AuNP clusters form at the presence of CBD in the target sample. The LED serves as light source to excite the floating AuNPs and the photodetector converts the light into electronic signals.

### 2.3 Pre-Incubation and Centrifugation Enhanced Detection

As discovered in our previous work^21^, the sample preparation methods, particularly incubation and centrifugation, have a significant impact on sensing accuracy and detection time of the AuNP sensors. Here we have compared two methods for CBD detection, i.e. incubation only (schematic and results in Figures S1 and S2) and a new approach termed ICED, namely preanalytical Incubation of CBD with binder-functionalized AuNPs, <u>C</u>entrifugation, and <u>E</u>lectronic <u>D</u>etection. For the incubation-only approach, 80 nm AuNPs coated with purified CA14 and DB21 were incubated with the CBD molecules for 4 hours (Figure 2a). Then, the top 5 µL liquid was pipette-loaded in a custom-made PDMS plate reader and spectroscopically measured using a UV-visible spectrometer coupled to an upright microscope. A significant color contrast was observed between 100 nM CBD and the negative control (NC, i.e. with no CBD but only buffer) (Figure 2a). To quantify the CBD detection, we further extracted the extinction intensity at the peak wavelength (∼560 nm) for 80 nm AuNPs and plotted it against CBD concentrations. The incubation-based sensing yielded a LOD of ∼ 0.7 nM (Figure 2h), comparable to the sandwich ELISA based detection method applied previously with the same set of co-binding CID system (LOD of ∼0.8 nM for ELISA) ^15^. The incubation-based system proved the feasibility of detecting small molecules such as CBD with a detection time similar to ELISA (typically around 5 hours) but still much faster than traditional mass-spec systems.

**Figure 2.**
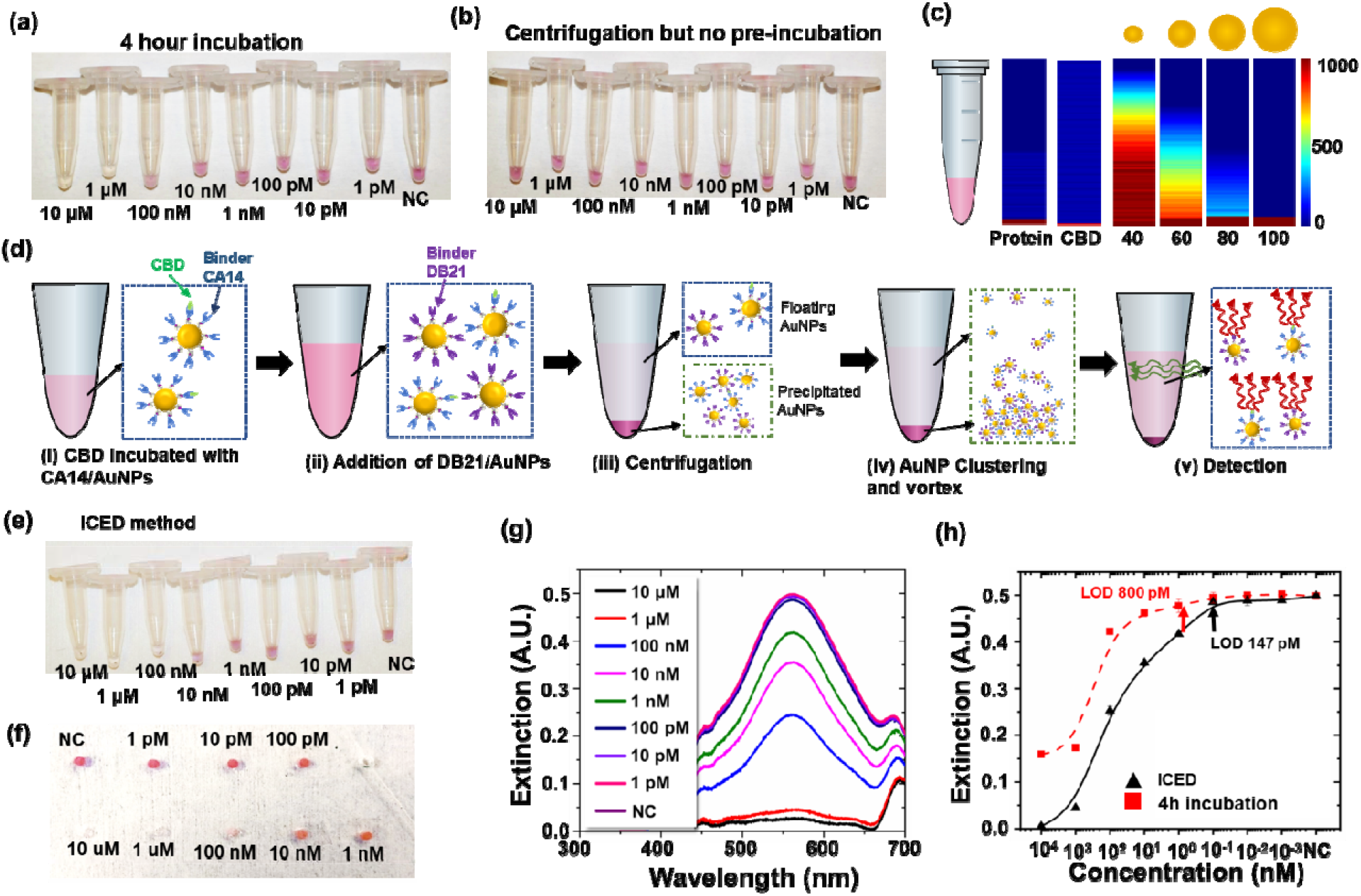
ICED detection of CBD molecule in PBS buffer. (a-b) Optical images of microcentrifuge tubes of CBD detection without preincubating CBD sample on AuNPs: (a) CBD mixed with CA14 and DB21-functionalized AuNPs, and then 4-hour incubation was applied. (b) CBD mixed with functionalized AuNPs, and then centrifuged, incubated for 20 min and vortexed. (c) Comparison of simulated molecule and AuNP distributions across liquid height after centrifugation. (d) Schematic showing process flow of ICED detection of CBD molecule. Here CA14-functionalized AuNPs was preincubated with to-be-tested CBD samples prior to mixing with DB21-functionalized AuNPs for sensing. (e-f) Optical images of CBD detection using ICED method: (e) microcentrifuge tubes; (f) PDMS plate image of extracted top liquid from Figure e. (g) Extinction spectra of AuNPs from PDMS plate in Figure f. (h) Extinction peak values (at 559 nm) for ICED method (extracted from plot Figure g, Black triangle) and after 4-hour incubation (red squares).

To further improve the assay sensitivity and reduce the assay time, we theoretically studied the impact of AuNP sedimentation process on sensing. In such an AuNP-based sensing system, two critical parameters governing the assay time and performance are aggregation time constant τ_α_ and sedimentation time constant τ_α_. In determining τ_α_, we employed a simplified version of Smoluchowski’s coagulation equation^27^ to estimate an empirical parameter P, defined as the probability of a binding event resulting from each molecular collision, and found the best estimated P value was ∼1. This indicated that the multivalence of the conjugated AuNPs increased the potential binding affinity compared to mono-binding process^22, 23^, significantly different from traditional surface-based detection such as ELISA ^28, 29^. This also implied that ELISA-determined binding constants were non-ideal to precisely predict the effective affinity observed in our solution-based multivalent sensing system. Additionally, τ_α_ can also be reduced significantly by increasing the AuNP concentration, which could be achieved by applying centrifugation to localize the AuNPs at the tube bottom. Further, we calculated τ_s_ = z/(s • g) using Mason-Weaver equation^30, 31^, where *z* is the precipitation path (for example, the height of colloid liquid), *g* is the gravitation constant, and α is the sedimentation coefficient dependent on the physical properties of AuNPs and buffers^30^. For a liquid height of z∼3.5 mm (approximated for 16 µL liquid in a 0.5 ml Eppendorf tube), we found the τ_α_ reduced from 26 hours for 80 nm AuNP to 20 mins for 800 nm clusters, respectively. This suggested that larger AuNP clusters, formed during CBD to nano-binder binding, would precipitate rather quickly. Assisted by simulation, we could also visualize that 80 nm AuNPs had a very narrow equilibrium gradient distribution in micro-molar concentration^30, 31^, more specifically that most of the dimers and trimers resided at the tube bottom. Given that the signals were only collected from the top-layer solution, only floating monomers modulated the observed optical extinction intensity, while the sedimented AuNP dimers or oligomers would not contribute to the solution signals.

The theoretical analysis pointed out that increasing the AuNP concentration and decreasing the liquid height, i.e. volume, would enable us to detect CBD faster. However, too high AuNP concertation would have resulted in saturation of optical extinction at lower CBD concentration, thus lowering detectability. On the other hand, it is challenging to reliably collect the signals only from free-floating AuNPs in the top liquid of a very small liquid volume. We applied an centrifugation approach to accelerate the detection (schematic in Figure S3), similar to the method we employed in rapid Ebola and SARS-CoV-2 antigen sensing^21^. However, the incubation time (1 to 20 min) used for protein sensing was found insufficient here to produce a visible color change except at the highest (10 µM) CBD concentration (Figure 2b). To investigate this phenomenon, we used Stokes centrifugal force equation to calculate the spatial distribution of particles (Figures 2c and S4) under centrifugal force^32^ for CBD, proteins and AuNPs of different sizes from 40 to 100 nm. The simulation showed that this centrifugation step, although effectively concentrating the AuNPs at the bottom of the tube within about 2 minutes, is incapable of concentrating CBD. Still broadly distributed within the tubes, the amount of CBD molecules at the tube bottom was rather limited, and therefore the CBD-induced AuNP clustering process was slow and could be mainly CBD diffusion-limited.

To circumvent this slow reaction issue, in our new ICED approach we developed a strategy to enhance CBD concentration for improved detection. To do so, we opted to use the higher affinity binder CA14 from the CA14-DB21 pair (6 µM for CA14 compared to 56 µM for DB21) as a CBD carrier to transport target CBD molecules to the reaction zone at the tube bottom. More specifically, we pre-incubated the CA14 coated AuNPs with CBD for 2 hours, followed by mixing with the DB21 coated AuNPs and centrifugation (Figure 2d, and more data in Figures S5 and S6). This nano-binder/AuNP-mediated pre-incubation of CBD (Figure 2e) evidently improved the detection of CBD (Figure 2e). The concentration-dependent CBD sensing results were further quantified by extracting the top 5 uL in a PDMS plate (Figure 2f), examining their optical extinction (Figure 2g) and recording the extinction peaking values at the AuNP resonance (560 nm) (Figure 2h). Clearly, the incubation-based (red line, Figure 2h) and ICED methods (black line, Figure 2h) displayed similar concentration-dependent signal moduclation; however, the LOD for the ICED method (∼144 pM) was about 5 times better than the incubation-based detection (∼700 pM) and the traditional Sandwich ELISA with the same reagents (∼800 pM)^11^.

### 2.4 Modular Analytic Model for Sensor Optimization

The underlying detection mechanism involves multidisciplinary studies of analyte-ligand binding, plasmonic AuNP effects, and AuNP aggregation and sedimentation behavior. The molecular sensing process involves complex biochemical binding between the target molecule to binder pairs and physical processes for AuNP clustering and sedimentation, and therefore a number of factors, including multivalent molecular binding, sedimentation time, aggregation, etc., can affect the assay preformation. In order to better verify the working mechanism, we tested CBD with 0.036 nM AuNPs, extracted the top-level liquid, and diluted for nanoparticle tracking analysis (NTA) measurement (supplementary Figure S7a). From the NC sample, the AuNP size distribution (average 80 nm, 3-sigma deviation 20 nm) was consistent with expectation, considering the coating of Streptavidin and nano-binders on the AuNPs. Such NTA analysis served to verify that the top liquid only contained monomers of the functionalized AuNPs. In addition, we also extracted 2 uL liquid of all the samples from the tube bottom, dried the liquid, and imaged the samples by transmission electron microscopy (TEM) (supplementary Figures S7b and Figure S8). Clearly, only AuNP monomers were observed for the NC sample, evidently showing minimal non-specific AuNP clustering, but AuNP clusters of different sizes were only found for CBD concentrations from 1 pM to 100 nM, providing evidence of AuNP cluster formation in sensing.

In order to formulate a model to better understand the complex chemical reactions and fluidic dynamics involving CBD molecules, nano-binders, and AuNPs, we created a modular analytic model (Figure 3). Briefly, we started with a dynamic model for the reaction kinetics in combination with Smoluchowski’s coagulation equation and utilized two different coagulation models, i.e. Mason – Weaver equation for gravitational sedimentation and Stokes’ equation for centrifugal sedimentation, to parameterize the initial conditions as well as to predict the aggregate precipitation. Additionally, we identified two parametric factors that correlate to the enhancement effect of our system, i.e., NPf signifies the multivalence effect of the nanoparticle sensor (proportional to the surface area or d^2^) while Cf signifies the nanoparticle concentration effect by centrifugation. In addition, the pre-incubation was introduced to favor the chemical reaction of Capture nano-binder (CA14) and CBD to form the CA14-CBD complex on AuNPs (Figure 3a). We found that Npf = 50 and Cf = 100 produced the best experimental fitting for our assay using 80 nm AuNPs (Figure S9a). The centrifugation effect (Figure S9b) was visualized that the incubation-based assay (red curve, Cf = 0) had a much worse signal contrast in optical extinction at the highest analyte concentrations (10 μM) compared to that with ICED approach (black curve Cf=100). Further, we also simulated the impact of the inherent antigen-antibody binding affinity on the assay performance (Figure S9c), where two different nano-binder systems CA14-DB21 (Kd= 56 nM) and CA14-DB18 (Kd = 560 nM) were used. Interestingly, despite that DB18 has one order of magnitude lower binding affinity, its predicted sensing signal contrast, i.e. the optical extinction intensity difference at the highest CBD concentration (10 µM) from the NC signal, was only moderately worse (at 0.18) compared to that from DB21 (0.08). This is partially attributed to the fact that DB18 has a higher dissociation constant (560 nM compared to 56 nM) but a comparable association constant (ka)^10^. The molecular dissociation is partly compensated by the use of multivalent in-solution AuNPs coated with many binders, which effectively promotes stronger binding. Further, the precipitation-based readout was much less reversible and more favorable compared to monolayer analyte binding in a conventional ELISA assay. Lastly, a good agreement between our model fitting for the 80 nm AuNP and the experimental results proved the validity of our analytical model to guide sensor design and optimization in similar affinity-based assays, from sensing of small molecules to proteins and even to nucleic acids. In addition, coupling modeling with experimental analysis will also serve to better design and screen the performance of designed synthetic nanobinders.

**Figure 3.**
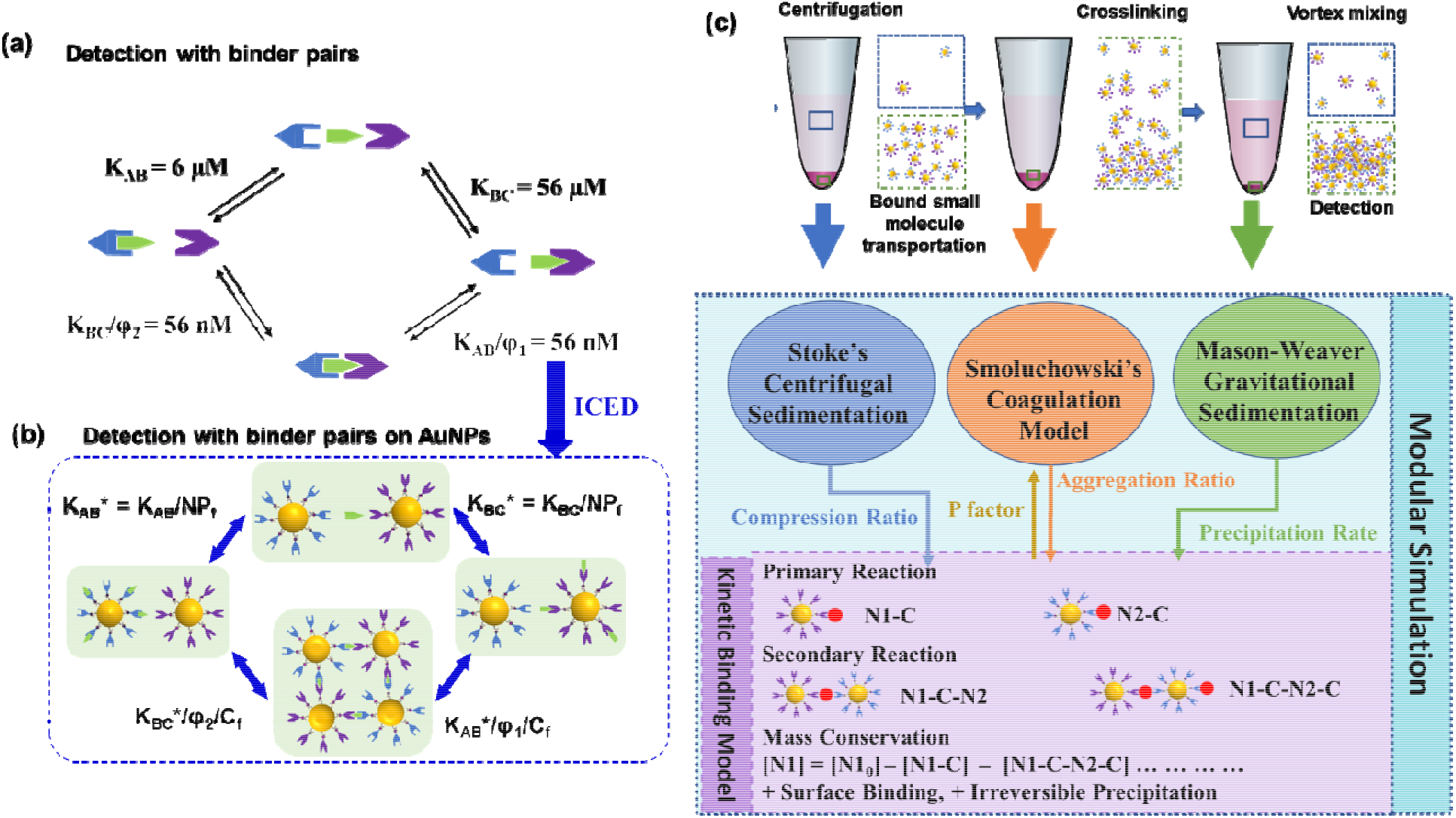
Modular analytic model for ICED assay. (a-b) Schematics showing the enhancement in effective binding affinity with AuNPs: (a) a typical sandwich assay with an effective binding affinity of 56 nM using a pair of anchor binder (CBD binding affinity of 6 µM) and dimerization binder (CBD binding affinity of 56 µM); (b) the use of multivalent AuNPs effectively improves the binding affinity by a factor of C_f_. (c) Model framework showing modular simulation strategy comprising of individual physical models used to simulate reaction mechanism for multivalent AuNPs aggregation and sedimentation based bio-sensing system.

### 2.5 Assay Optimization

### 2.5.1 Impact of Nanoparticle Sizes on Sensing

To evaluate the role nanoparticle size plays in CBD detection, we employed the CBD ICED assay with different AuNP sizes (40, 60, 80 and 100 nm) (Figure 4a-d, more data in supplementary Figures S10, S11, S12, and S13). Clearly, the color of 1 uM CBD concentration could be easily differentiated from the NC sample for all AuNP sizes. However, further analysis of the extinction signals shows that 80 and 100 nm AuNPs were better in dynamic range and LOD (Figure 4b). Additionally, the impact of AuNP size was also evaluated by our modelling (Figure 4c). Clearly, the signal intensity contrast was much higher for larger nanoparticles (0.5 for 80 nm AuNPs and 0.45 for 100 nm AuNPs) than for smaller ones (0.3 for 40 nm AuNPs and 0.35 for 60 nm AuNPs, respectively). This can be attributed to two factors, i.e. effective sedimentation described previously (section 2.3) and effective concentration [NP]_e._ Here [NP]_e_ of a multivalent nanoparticle sensor is defined as the product of the concentration of the nanoparticle[*NP*] and the amount of surface ligands per *NP* [*S*] following *[NP]*_*e*_ ∝ [*NP*][*S*].

**Figure 4.**
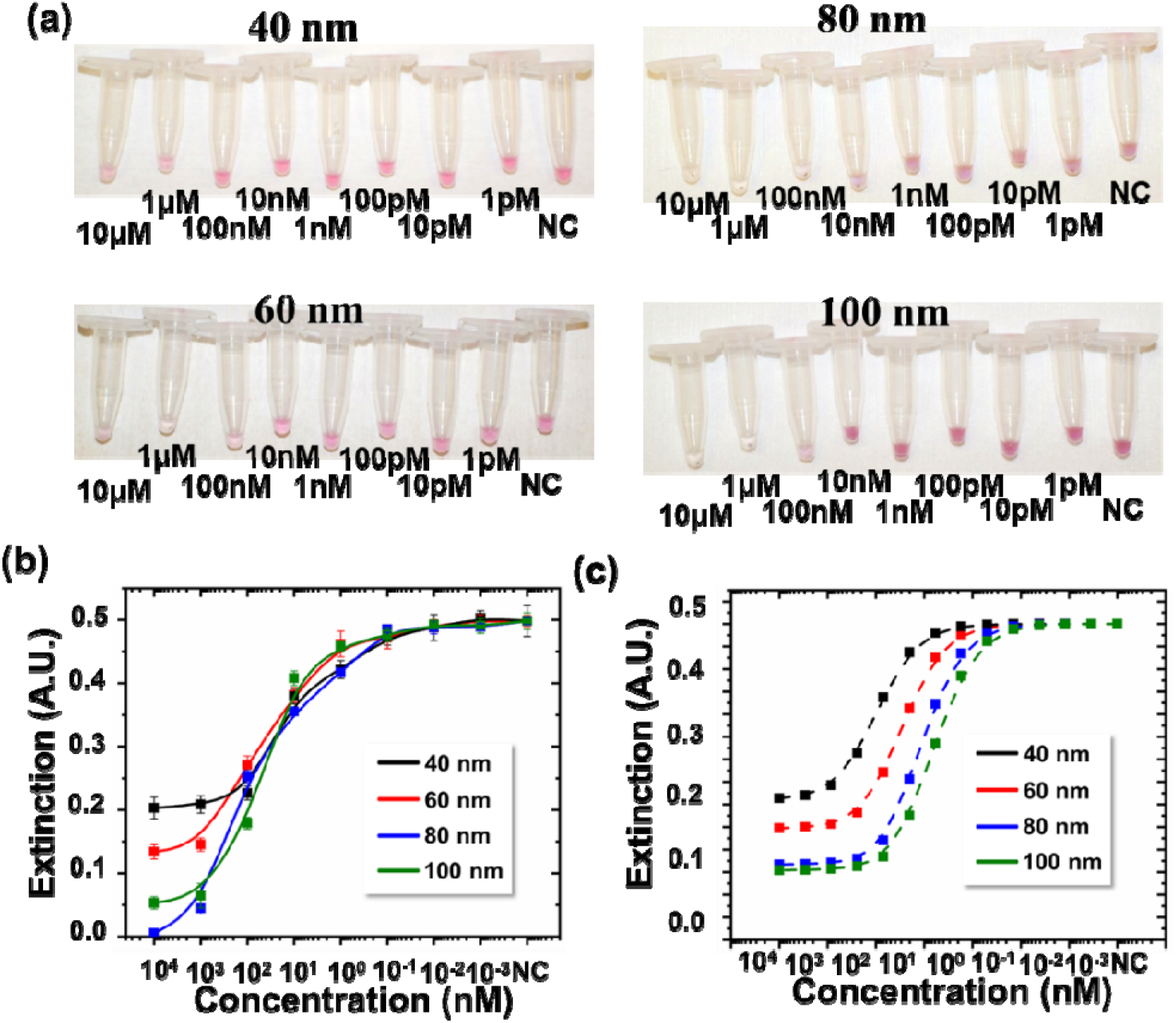
Experimental and simulation analyses of nanoparticle size effect on CBD detection. (a) Optical images of CBD detection in PBS for AuNP sizes of 40 nm, 60 nm, 80 nm and 100 nm. The pictures were taken after all reactions. Initial and intermediate-stage images were provided in supplementary figures. (b) Extracted optical extinction peak values for CBD detection in PBS for different nanoparticle sizes. (c) Simulated extinction peak values for CBD detection for dimerization system CA14-CBD-DB21.

Fundamentally, our sensor essentially monitors the localized surface plasmon resonance (LSPR) extinction of free-floating AuNPs. The LSPR extinction is dependent on the AuNP concentration [*NP*] and diameter *d* roughly following σ-_*ext*_ ∝ [*NP*]*d*^329^, and therefore the AuNP concentration in our test was lower at larger sizes following 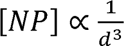 considering equal extinctions of AuNPs for different sizes in our experiments. Further considering that [*S*] is also related to the surface area and hence diameter of the particles as [*SJ*] ∝ *d*^2^, we obtain [*NP*] ∝ 1/*d*. Therefore, a higher [*NP*]_*e*_ for relatively small AuNPs under our experiment conditions was favored to promote reaction and signal readout. Indeed, too large nanoparticles did not perform well due to their lower particle concentrations.

However, on the other hand, sedimentation time also increases with smaller AuNPs. This is particularly true for detection by just incubation, where typically a few hours are needed for effective AuNP clustering and subsequent precipitation. In comparison, for ICED method, the use of centrifugation effectively concentrates the AuNPs to promote faster reaction. For example, we have performed Stokes’ centrifugal simulation (Figure S4) to show that a 2-min centrifugation at 1200g could concentrate 80 nm AuNPs by a factor of ∼50 but only by ∼10 for 40 nm AuNPs. Clearly, smaller nanoparticles still occupies a larger volume even after being driven by centrifugal forces. This results in a longer precipitation path and a longer sedimentation time, diminishing their advantage in higher initial concentration and faster molecular reaction. In general, we observed a good overall agreement on the effect of nanoparticle size (Figure 4b-c) between experiments and model prediction. Experimentally, 80 nm AuNPs performed better than 100 nm AuNPs and thus were chosen as the primary design for CBD sensing.

#### 2.5.2 Nano-binder impact

Further, two nano-binder pairs CA14/DB21 (Kd = 56 nM when sandwiching CBD) and CA14/DB18 (Kd = 560 nM for CBD) were compared for their sensing performance. Even though the overall binding affinity of DB21 was 10 times higher compared to using DB18, its resultant LOD was only 5 times better than that of DB18 (Figure 5a, Figure S5-6 and Figure S14). This could be attributed to a high association constant (ka) of DB18, which is comparable to that of DB21, and the induced AuNP aggregation. As predicted in our model (Figure S9c) and verified by our experiments, we suspect that binders with comparable binding affinity but with high association (ka) and dissociation (kd) constants would be beneficial to produce a better LOD, because high dissociation would be partially suppressed by the presence of multivalent binding sites on the AuNPs, which essentially suppresses dissociation and promotes AuNP clustering. Clearly, this example suggests the complexity of the sensing system and the importance of combination of analytic modeling and experimentation in studying the reaction mechanism. Further, it may also shed light onto the designs and selection of antibodies and nano-binders, where both the binding affinity and the association constants may play important roles.

**Figure 5.**
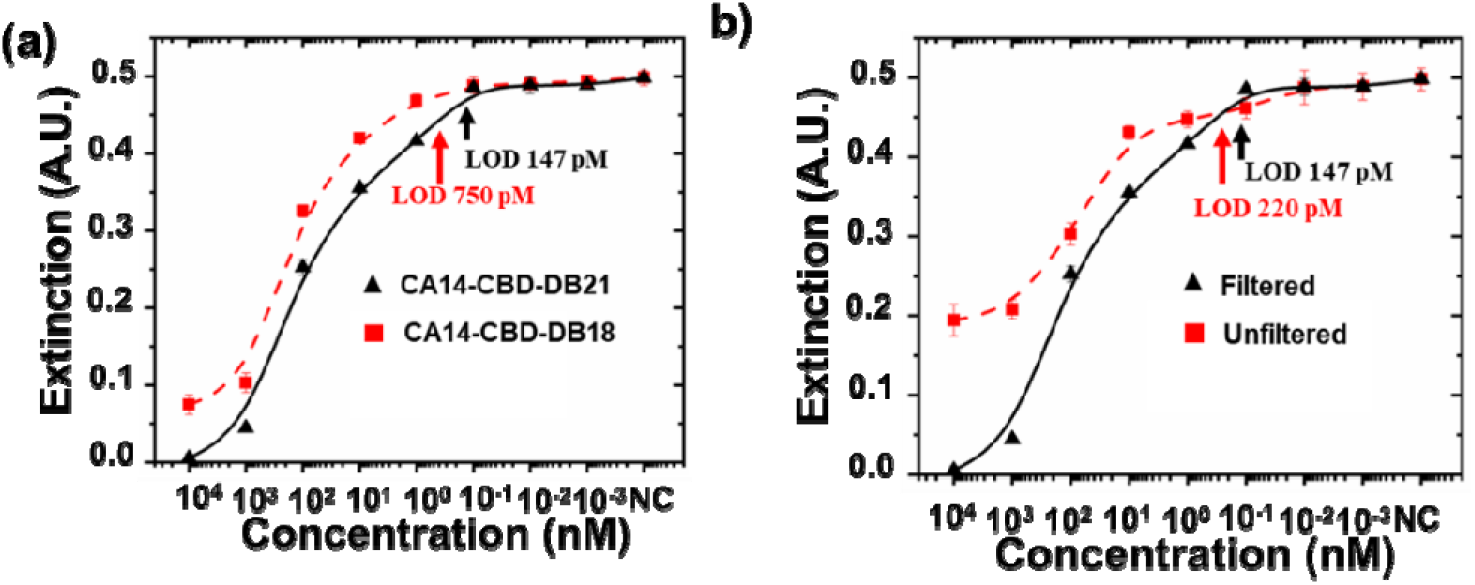
Impact of nano-binder and reagent filtration on CBD detection. (a) Extracted optical extinction peak for CBD detection in PBS for dimerization system CA14-CBD-DB21 (Black triangle with solid line) and CA14-CBD-DB21 (red square with dashed line). (b) Extracted optical extinction peak values for CBD detection in PBS for filtered (black triangle with solid line) and unfiltered (red square with dashed line) assay.

#### 2.5.3 Unfiltered Assay Development for lower coast remote area application

The coupled experimental and modeling analysis proved that our design strategy was successful in small-molecule analyte detection in body fluids with greatly improved sensitivity compared to ELISA. To further explore the feasibility with even simpler sensing scheme, we tested our system with unfiltered assays, i.e., mixing the nano-binder solutions with AuNPs without any post-conjugation purification. In particular, 80 nm AuNPs was mixed with nano-binders at a molar ratio of 1:1,280 (based on the estimated number of binders per each AuNP) and incubated for 30 min, and then used to detect CBD following the ICED method. Despite worse performance (LOD of 750 pM) compared to using filtered AuNPs (LOD ∼147 pM, Figure 5b, and additional data in Figures S15-19), possibly due to the presence of partially conjugated AuNPs and excessive unbound nano-binders (Figure S20 and Table S1), this approach was much simpler, because it not only skipped the purification process but also reduced the AuNP consumption by half and reduced nano-binder consumption by 10-15 times (excess amount was needed for conjugation but then lost during filtration, Table S1). Additionally, this method eliminates the necessity for a high-speed centrifuge. This sample preparation method could be particularly of interest in applications where portability, cost, and accessibility are of great importance.

### 2.6 Detection of CBD in Urine and Saliva using Portable Electronic Detection System

To further facilitate portable CBD detection, we demonstrated a portable electronic detection (PED) system comprising a tandem LED and photodiode system (Figure 6a). The LED emit at a wavelength matching that of the 80 nm AuNP extinction peak, and the light passing through the upper-level liquid was collected by a photodiode, where electrical signals was produced and adjusted using serially connected load resistor. For demonstration, a snug-fit microcentrifuge holder was 3D-printed, where windows were open to define the optical path along the upper portion of the liquid. The PED system was validated to produce a large dynamic range so that large variations in CBD concentration can be measured without saturation. In practice, we measured CBD in 5% urine and saliva, in both testing tubes (Figure 6b-c and Figure S21-22) and PDMS well plate. The optical measurement results showed a LOD of ∼165 pM and ∼198 pM for CBD spiked in urine and saliva (Figure 6d-e), as well as a broad dynamic range (5 logs) and excellent specificity (∼3 logs more concentrated THC is required to equitable signal level) against THC (Figure 6d-e and additional data in Figures S23 and S24). In comparison, the PED system further improved the LOD to ∼88.5 and ∼97.5 pM for urine and saliva, respectively, or ∼8 times better than tested by ELISA ^18^(Table S2). This PED-enhanced performance can be attributed to lower 3-errors generated during electronic measurements^21^. With a small footprint, low-cost readout device (a few dollars) and low reagent cost (estimated <$0.1 each test)^21^, reliable yet simple operation, and a potential for automatic data collection, the PED-based small molecule detection method has great potential for affordable high-throughput applications.

**Figure 6.**
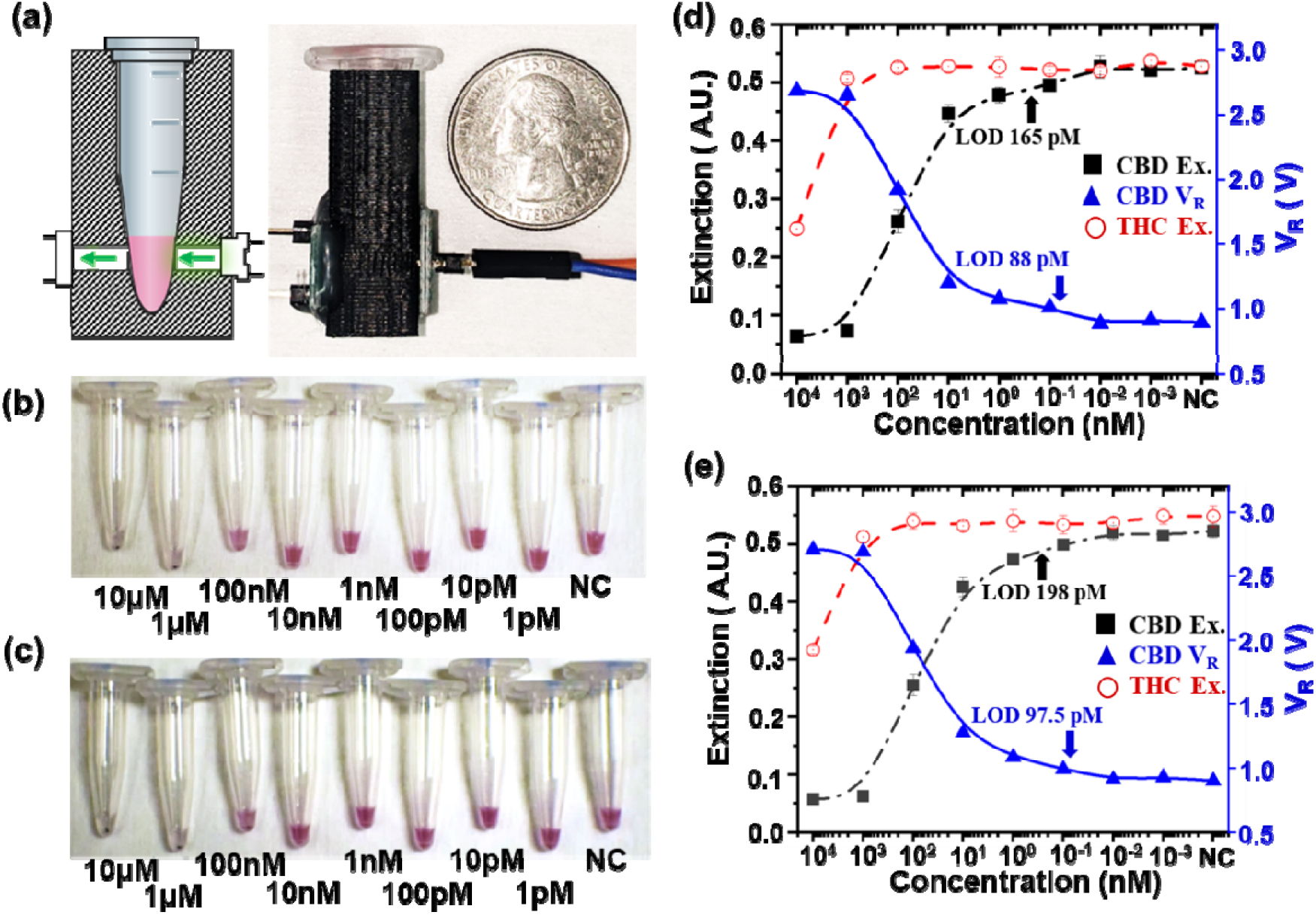
Rapid and portable electronic detection of CBD molecule in urine and saliva. (a) Schematic and optical image of PED system, consisting mainly of a LED circuit, a photodiode circuit, and a 3D printed Eppendorf tube holder. (b) Optical image of detecting CBD spiked in urine. (c) Optical image of detecting CBD spiked in saliva. (d-e) CBD and THC sensing in: (d) 5% urine and (e) 5% saliva. Black squares and fitted dash-dot line: optical extinction peak values (559 nm) in CBD sensing were extracted from optical spectra (Figure S21 e and S22 e) and plotted against CBD concentration. Red open-circle and dash line: optical extinction peak values (559 nm) in THC sensing were extracted from optical spectra (Figure S23 e and S24 e) and plotted against THC concentration. Blue triangles and solid line: Electronic voltage signals measured by PED in detecting CBD from the tube samples shown in Figure 6b and 6c.

## 3. Conclusions

We have demonstrated a generalized rapid assay design to achieve sensitive detection of small molecules, using CBD (Mw 314.47 g/mol), an important molecular target in detection of drug misuse^33^, as an example. Here, a multi-leveled sensitivity-enhancing scheme has been innovated. First, a pair of nano-binder was used to form a stable sandwich with CBD at an improved affinity (56 nM compared to 6 µM and 56 µM individually). Secondly, multivalent AuNP sensors were used, each hosting hundreds of binding sites, thus greatly improving the effective binding affinity. In addition, we introduced a new ICED (i.e. brief incubation followed by centrifugation prior to electronic detection), to attach CBD to one set of AuNP sensors. The centrifugation process transports CBD molecules to the reaction zones at the testing tube bottom, thus greatly localizing the CBD and boosting their concentration prior to detection for improved sensitivity while decreasing the detection time. In addition, the AuNP aggregation and sedimentation occurs at presence of targeted small molecules, enabling CBD-concentration-dependent optical extinction display and subsequent portable electronic readout, which does not collect signals from other background materials and minimizes the background noise. As a result, this new sensing method can overcome the traditional challenges of forming reliable readout signals from low-affinity small-molecule binders, instead achieving high sensitivity (<100 pM), large dynamic range (5 logs), and high specificity against THC in biological medium. Additionally, this assay format eliminates long incubation or cumbersome washing steps typically required for ELISA or surface-based detection and greatly decreased the footprint for readout, making it an ideal platform for affordable and accessible detection in resource limited regions.

## 4. Materials and Method

### Materials

Phosphate-buffered saline (PBS) was purchased from Fisher Scientific. Bovine serum albumin (BSA) and molecular biology grade glycerol were purchased from Sigma-Aldrich. Sylgard 184 silicone elastomer kit was purchased from Dow Chemical. DNase/RNase-free distilled water used in experiments was purchased from Fisher Scientific. The Streptavidin-functionalized AuNPs were purchased from Cytodignostics, dispersed in 20% v/v glycerol and 1 wt% BSA buffer.

### Nano-binder generation, Selection, Expression, and Biotinylation

In brief, rationally designed CDR sequences of different ratios of amino acids were used to generate a library of nano-binders. Anchor binders were selected after six rounds of biopanning using Biotin and Biotinylated-CBD-bound Streptavidin magnetic beads. Dimerization binders were selected after four rounds of biopanning using CBD-free and bound CA14 (anchor binder selected in the previous step) as the negative and positive control. All nano-binders were C-terminal AviTagged before Biotinylation. Nano-binders bearing AviTag were biotinylated by BirA using a BirA-500 kit (Avidity).

### Preparation of Nano-binder surface-functionalized AuNP colloidal solution

AuNPs of different sizes can be used for experimentation and optimization, here the methodologies are provided using 80 nm AuNPs. Streptavidin-functionalized 80 nm-gold nanoparticles at 0.13 nM concentration were mixed with biotinylated nano-binders (capture binder and different dimerization binders) in excess, and incubated for 2 hours to ensure complete streptavidin-biotin conjugation. Next, centrifuge purification (accuSpin Micro 17, Thermo Fisher) was applied at 10,000 rpm for 10 mins and the top supernatant was discarded. This procedure was repeated twice to ensure high-quality purification. The concertation of purified AuNP colloidal solution was measured by Nanodrop 2000 (Thermo Fisher) and readjusted to 0.048 nM, which was chosen for optimal extinction-level during spectrometric measurement. The buffer used for mixing, dilution, purification, and subsequent readjustment contained 1 X PBS with 20 v% glycerol and 1 wt% BSA, prepared from 10 X PBS powder, ultrapure water, glycerol, and BSA. This buffer was used to ensure AuNP sensor stability and minimize nonspecific interactions.

### Preparation of CBD and THC analyte solution

CBD and THC solutions were serially diluted from 10 µM to 1 pM in detection media, which typically contained 1 X PBS with 20 v% glycerol and 1 wt% BSA. For detection in urine and saliva, this buffer was readjusted to have 1 X PBS with 20 v% glycerol, 1 wt% BSA, and 20 v% of either urine or saliva.

### ICED detection

CBD of different concertation was pre-incubated for 2 hours with capture antibody functionalized AuNP (CA14-AuNP) colloidal solution at 2:3 volume ratio. This was done to make sure maximum surface coverage with CBD-CA14-AuNP complex. Then chosen dimerization-binder-functionalized AuNP (DB21 /DB18 -AuNP) were mixed with the pre-incubated colloidal solution at a volume ratio of CBD: CA14-AuNP: DB (21 or 18)-AuNP at 2:3:3. After mixing, the solution was centrifuged at 3,500 rpm (1,200×g) for 1 minute. After incubation at a chosen time (typically 20 mins), this colloidal solution was vortexed mildly at 800 rpm for 5 seconds.

### PDMS well plate fabrication

PDMS well plate was created from Sylgard-184 elastomer. As detailed in our previous work, the PDMS was cured in a plain petri dish with a thickness of 2.5 mm. The PDMS film was cut into desired size and drilled 2 mm diameter holes with biopsy punch. Subsequently it was treated with oxygen plasma, and bonded to a glass slide of desired size.

### Spectrometric measurement

The UV-visible spectra and dark-field imaging were performed using a customized optical system (Horiba), comprising an upright fluorescence microscope (Olympus BX53), a broadband 75W Xenon lamp (PowerArc), an imaging spectrometer system (Horiba iHR320, spectral resolution 0.15 nm), a low-noise CCD spectrometer (Horiba Syncerity), a vision camera, a variety of filter cubes, operation software, and a high-power computer. Light transmitted through PDMS well plate was collected by a 50×objective lens (NA=0.8). The focal plane was chosen at the well plate surface to display the best contrast at the hole edge.

### Electronic Measurement

A LED-photodiode PED system was designed with three key components: a LED light source, a photodiode, and a microcentrifuge tube holder. The centrifuge tube holder was 3D printed using ABSplus P430 thermoplastic. An 8.6 mm diameter recess was designed to snuggly fit a standard 0.5 mL Eppendorf tube. Holes of 2.8 mm diameter were open on two sides of the microcentrifuge tube holder to align a LED (597-3311-407NF, Dialight), the upper-level assay liquid, and a photodiode (SFH 2270R, Osram Opto Semiconductors). The LED was powered by two Duracell optimum AA batteries (3 V) through a serially connected 35 Ω resistor to set the LED operating point. The photodiode was reversely biased by three Duracell optimum AA batteries (4.5 V) and serially connected to a 7 MΩ load resistor. The photocurrent that responds to intensity of light transmitted through the assay was converted to voltage through the 7 MΩ load resistor and measured with a portable multimeter (AstroAI AM33D).

### TEM sample preparation and imaging

To image AuNP precipitates, the supernatant was removed from the tube and 2 to 3 µL of AuNP sample colloid containing AuNP precipitates were left in the tube. The tube was then vortexed thoroughly. Samples of 2 µL were pipetted and coated on both sides of the oxygen plasma-treated Cu grid (Electron Microscopy Sciences, C flat, hole size 1.2 μm, hole spacing 1.3 μm). Then 30-second oxygen plasma was used for cleaning.

## Acknowledgements

This work was supported by grants from the U.S. National Institutes of Health (NIH) (1R35GM128918, R21DA051555, and R21DA051194) to L. Gu. C. Wang and M.A. Ikbal acknowledge partial support from National Science Foundation (NSF) under grant no. 1847324 and 2020464 and NIH under grant no. R21AI169098. The samples were characterized in the NanoFab cleanroom and Eyring Materials Center (EMC) at Arizona State University. Access to the NanoFab and EMC was supported, in part, by NSF grant no. 1542160.

## Notes

### Competing Interest Statement

The authors have declared no competing interest.

