## Supplementary document for "Picomolar-Level Sensing of Cannabidiol by Metal Nanoparticles Functionalized with Chemically Induced Dimerization Binders"

1. **Supplementary schemes and experimental data for AuNPs Based CBD Detection**

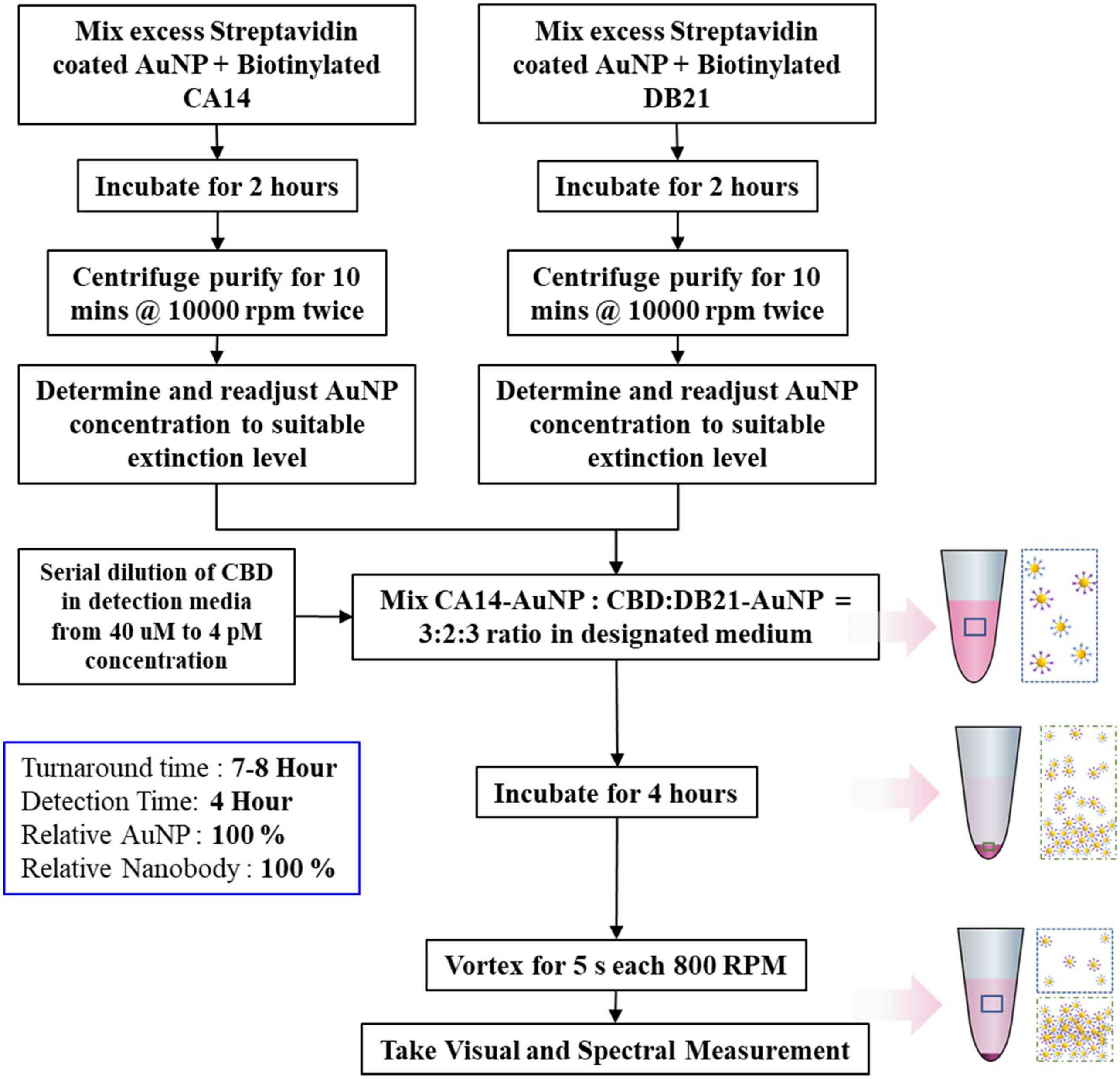


**Figure S1 Experimental Workflow of Incubation-Based CBD Detection Assay.**

Here CA14 and DB21 are examples of the nanobinder pairs for CBD detection.

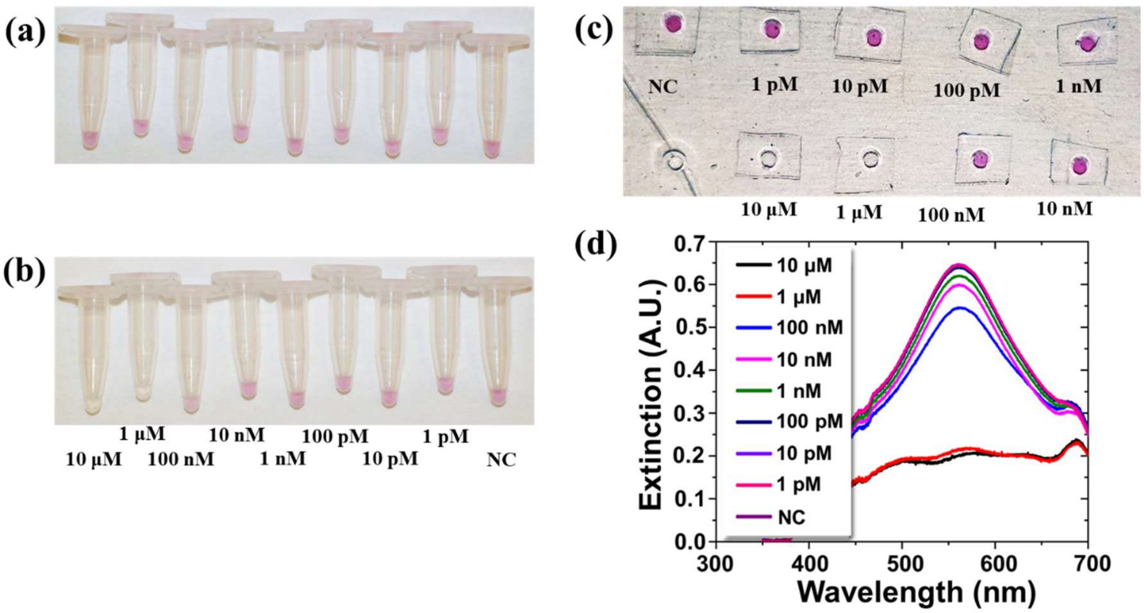


**Figure S2 Experimental Data for Incubation Based CBD Detection in 1X PBS with 80 nm Functionalized AuNPs**. (a-c) Optical images of microcentrifuge tubes (a) after mixing AuNP-CA14 sensor solutions with CBD molecules, (b) after mixing AuNP-CA14-CBD with DN21, incubation (4 hours) and vortex agitation. The CBD concentrations were labelled for the tubes. (c) Optical images of the upper-level liquid withdrawn from microcentrifugation tubes shown in Figure c loaded into a PDMS well plate. (d) Optical extinction measured from spectrometric analysis from the well plate plotted for different CBD concentrations. NC: negative control, where no CBD but only buffer was tested.

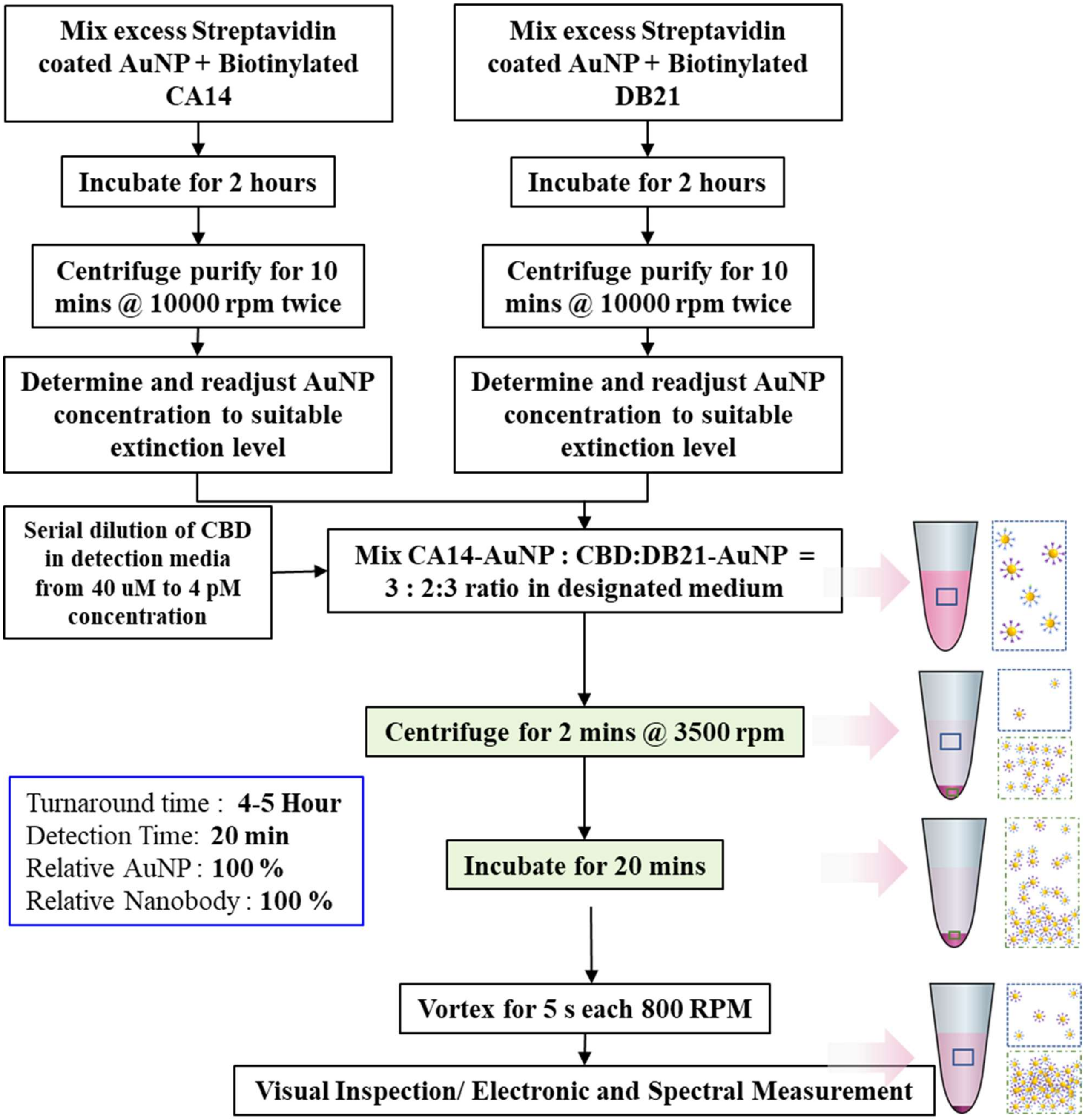


**Figure S3 Workflow Schematic of Centrifugation-Enhanced Method for CBD Detection.** Here a centrifugation step was introduced after mixing AuNPs with CBD and prior to incubation.

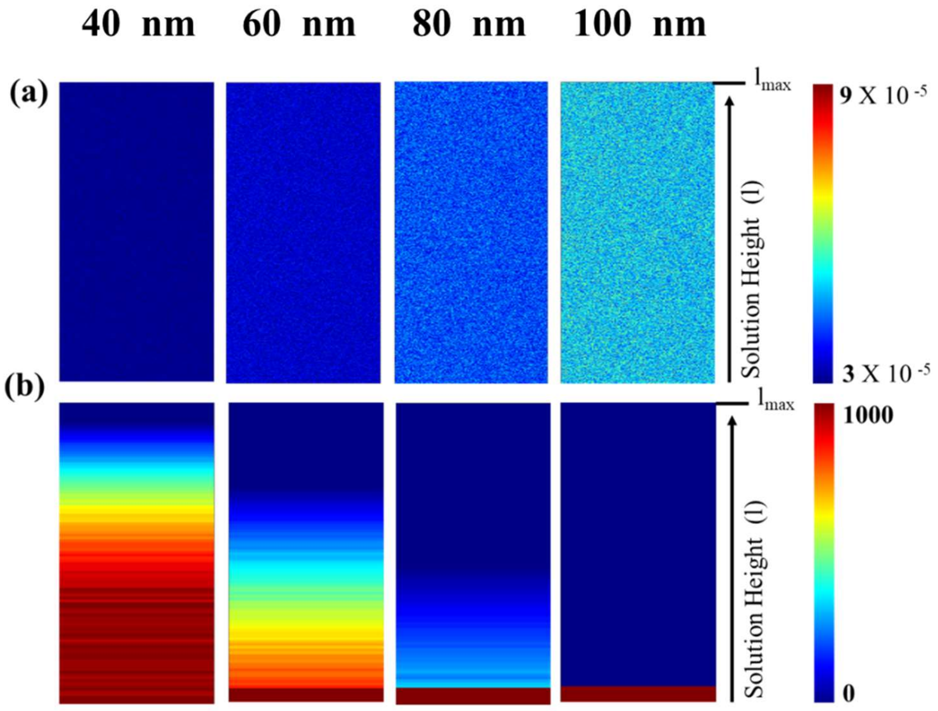


**Figure S4 Impacts of AuNP Sizes on Fluidic Properties in the Microcentrifuge Tubes.** (a) Sedimentation velocity of AuNPs. (b) Nanoparticle density distribution after centrifugation. The simulation was performed by solving Stokes equations. The lower boundary of the simulated area references the tube bottom.

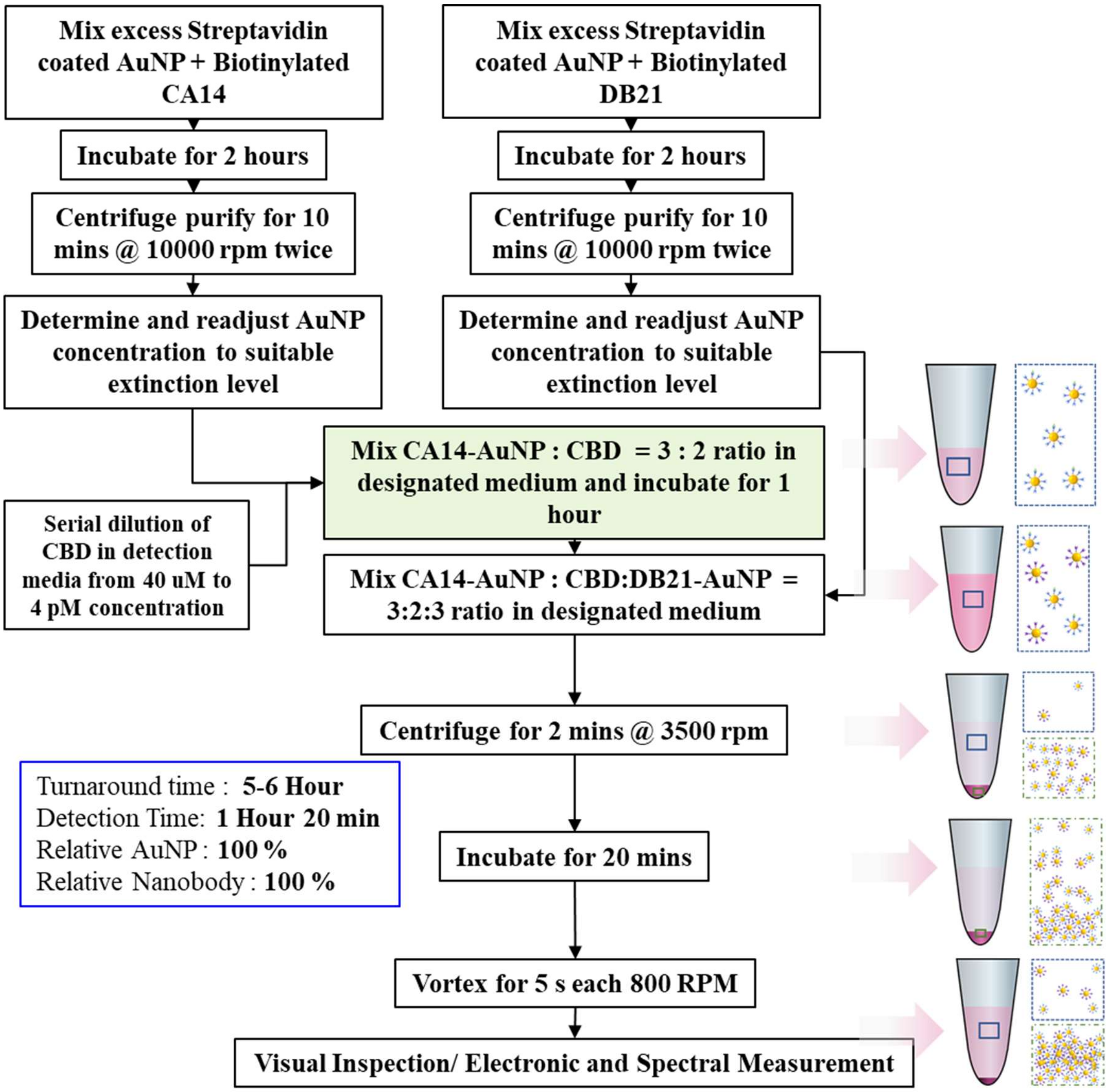


**Figure S5 Workflow Schematic of ICED Approach for CBD Detection**

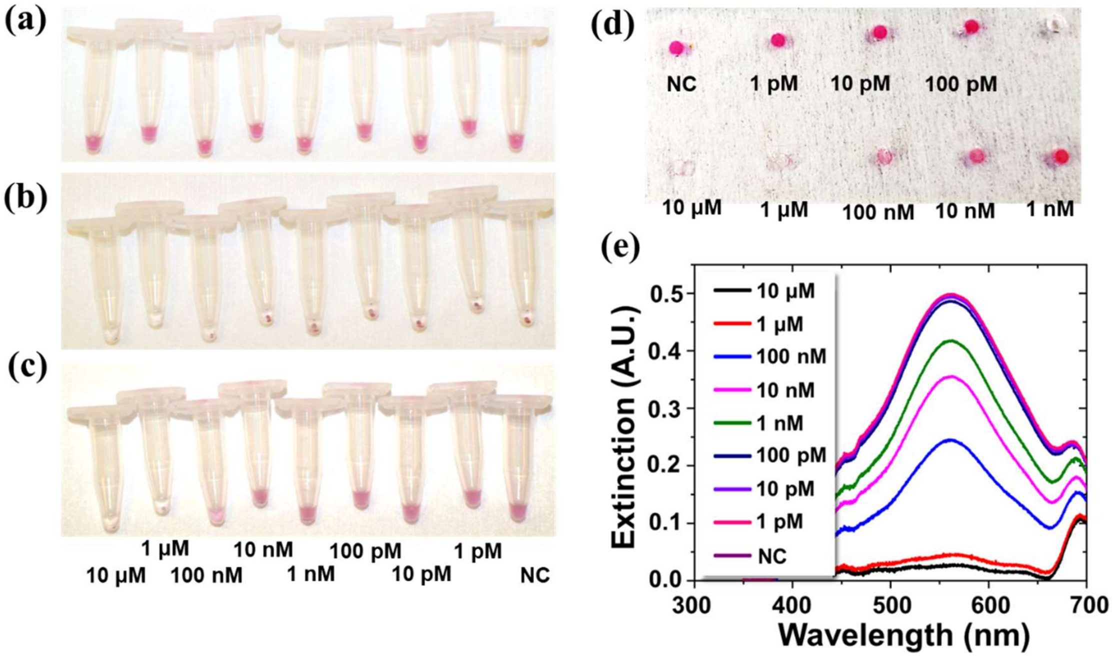


**Figure S6 Experimental Data for ICED-Based CBD Detection in 1X PBS with 80 nm Functionalized AuNPs**. (a-c) Optical images of microcentrifuge tubes (a) after mixing AuNP-CA14 sensor solutions with CBD molecules (Highlighted step in Figure S5), (b) immediately after mixing AuNP-CA14-CBD with DN21 and centrifugation, and (c) after incubation (20 min) and vortex agitation. The CBD concentrations were labelled for the tubes. (d) Optical images of the upper-level liquid withdrawn from microcentrifugation tubes (shown in Figure c) and loaded into a PDMS well plate. (e) Optical extinction measured from spectrometric analysis from the well plate plotted for different CBD concentrations. NC: negative control, where no CBD but only buffer was tested.

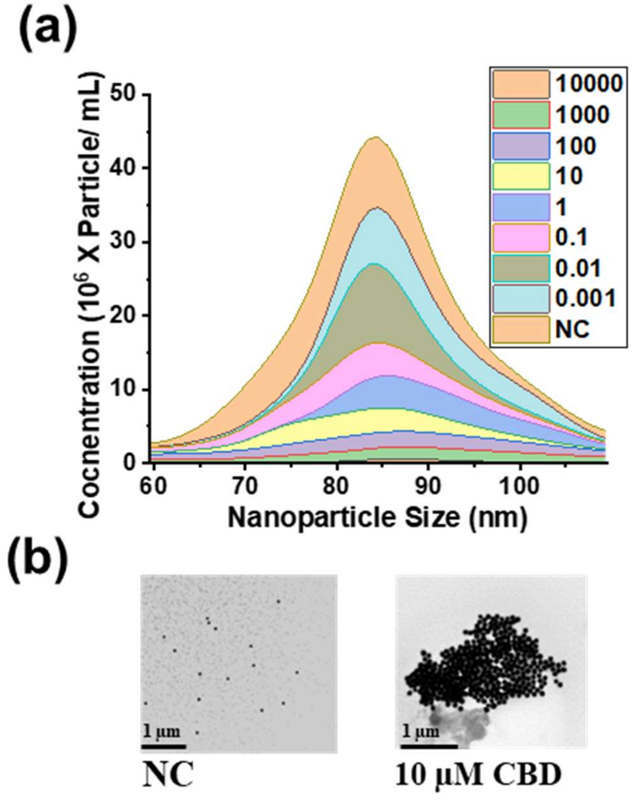


**Figure S7 Verification of monomer sampling and AuNP aggregate sedimentation.** (a) AuNP size and population data for CBD detection in PBS for 80 nm AuNP extracted from NTA analysis. (b) TEM image showing single AuNPs from NC sample and AuNP cluster form 10 uM CBD concertation sample.

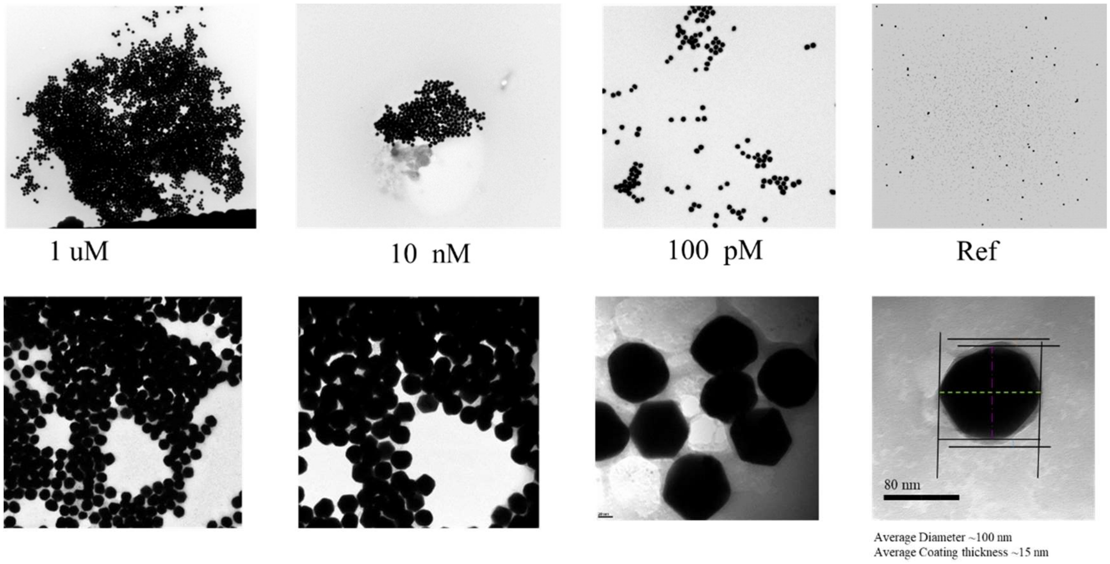


**Figure S8 Representative TEM Images of 80 nm AuNP Assay in Detecting 100 pM, 10 nM and 1 µM CBD , and NC Samples.**

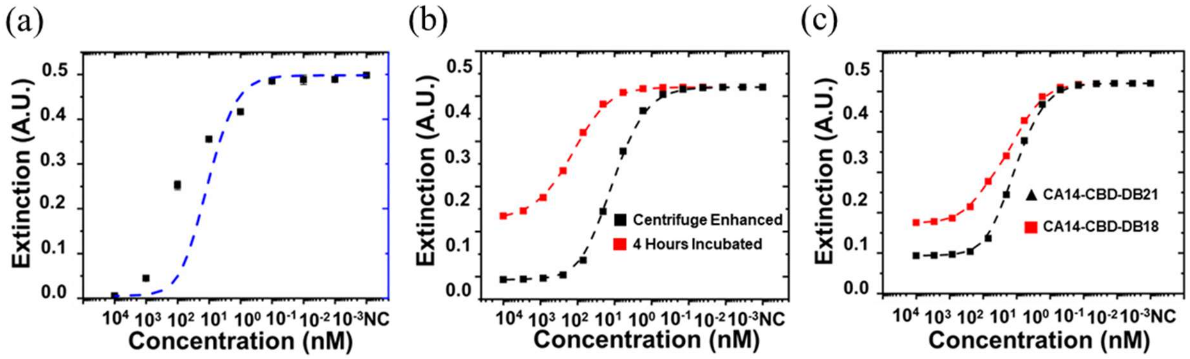


**Figure S9. Simulation results to compare the impacts of detection method and nanobinders.** (a) Simulated response curve (blue dash line) showing the centrifugation enhancement effect. The experimental data (Black dots) are consistent with simulation. (b) Simulated response comparing incubation-only (red) and centrifugation-enhanced (black) detection methods. (c) Simulated response showing the effect of different binding affinity arising from different dimerization binding system.

### 2. Study of the Impact of Nanoparticle Size on CBD Sensing Performance

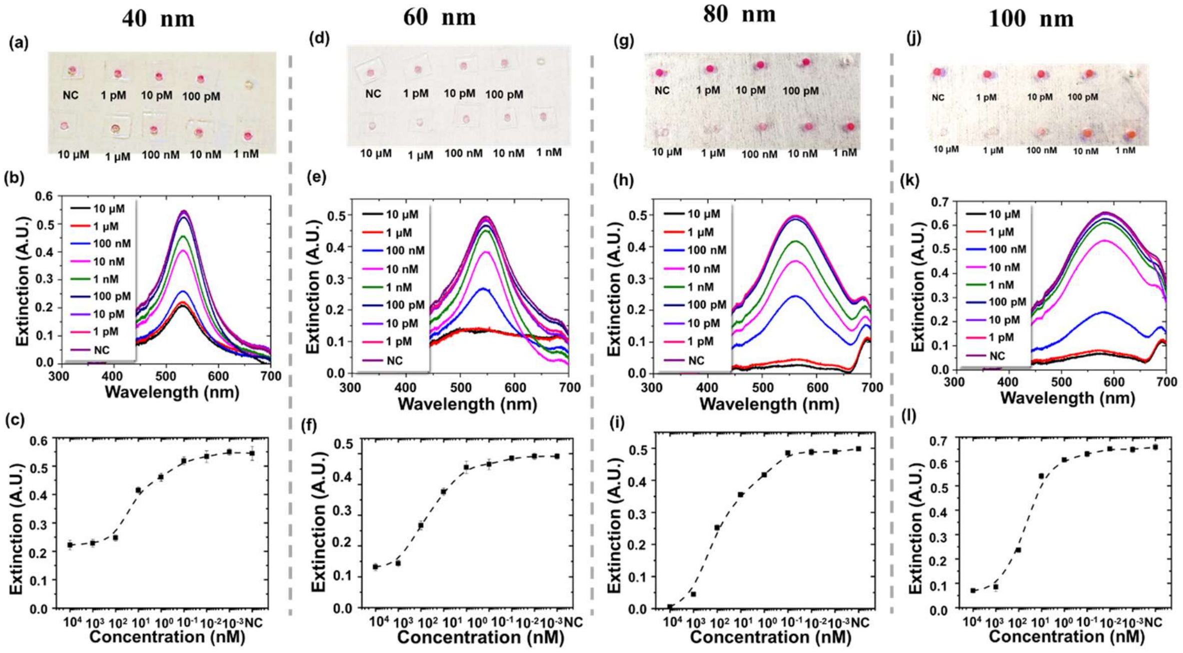


**Figure S10 Experimental analysis of the Impact of Nanoparticle Size on CBD Sensing Performance in 1 × PBS Buffer.** (a,d,g,j) Optical images of 40, 60, 80, and 100 nm AuNP assay, with the upper-level liquid samples extracted from the microcentrifuge tubes and loaded on a PDMS well plate for CBD detection. (b,e,h,k) Extinction spectra of samples shown in (a)-(d) panel, measured by a lab-based UV-visible spectrometer system. (c,f,i,l) Extinction peak values plotted for AuNP assay using different NP sizes in detecting CBD with concentration from 1 pM to 1 μM. Extinction values were derived from the peak extinction values in (b,e,h,k).

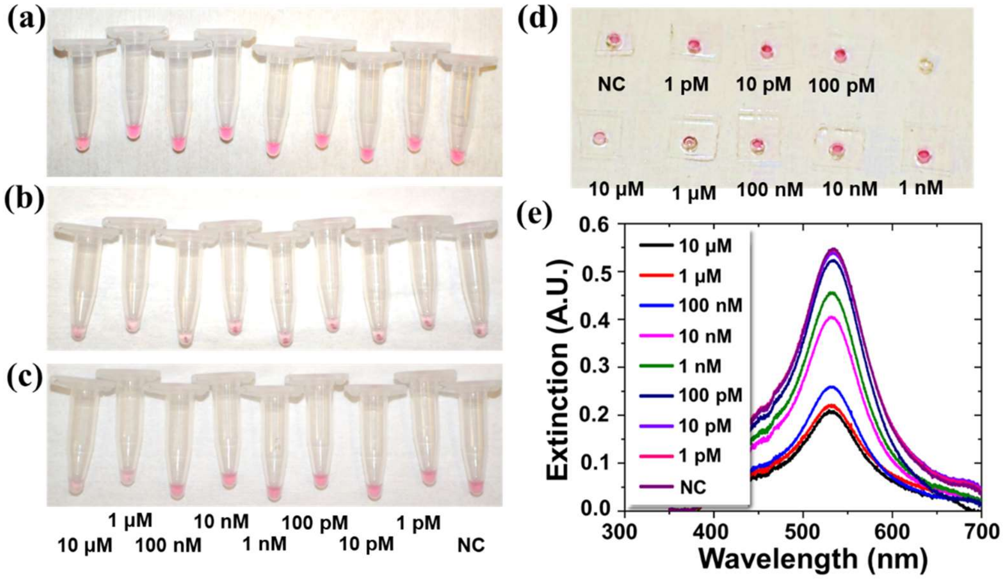


**Figure S11 Experimental results for ICED CBD Detection in 1X PBS with 40 nm AuNPs.** (a-c) Optical images of microcentrifuge tubes (a) after mixing AuNP-CA14 sensor solutions with CBD, molecules (Highlighted step in Figure S5), (b) immediately after mixing AuNP-CA14-CBD with DN21 and centrifugation, and (c) after incubation (20 min) and vortex agitation. (d) The upper-level liquid from (c) loaded into a PDMS well plate. (e) The spectrometric readout from the well plate.

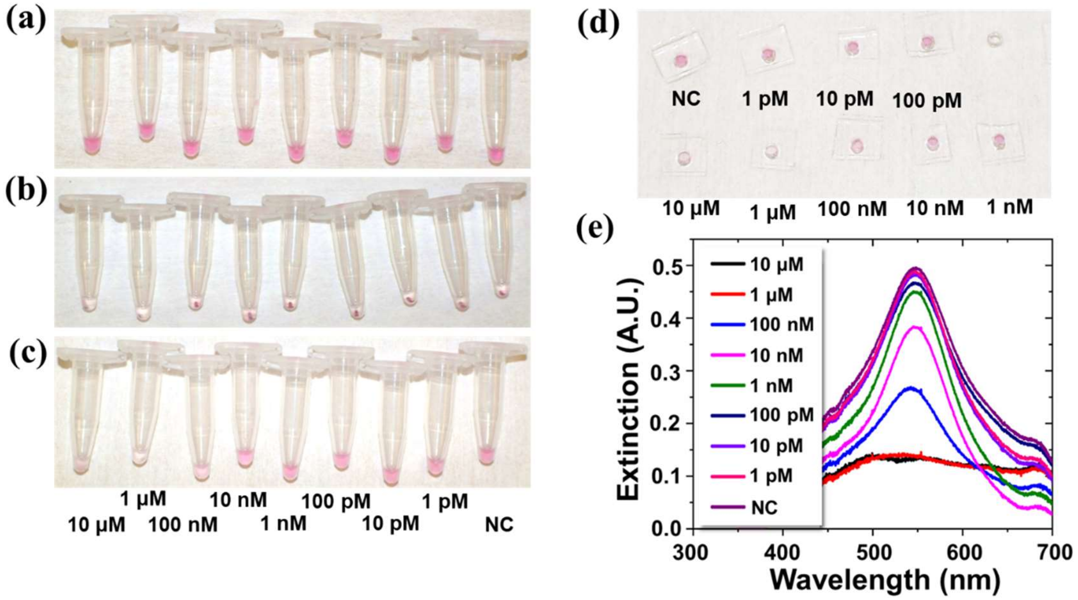


**Figure S12 Optical images for ICED CBD Detection in 1X PBS with 60 nm AuNPs.** (a-c) Optical images of microcentrifuge tubes (a) after mixing AuNP-CA14 sensor solutions with CBD, molecules (Highlighted step in Figure S5), (b) immediately after mixing AuNP-CA14-CBD with DN21 and centrifugation, and (c) after incubation (20 min) and vortex-mixing. (d) The upper-level liquid from (c) loaded into a PDMS well plate. (e) The spectrometric readout from the well plate.

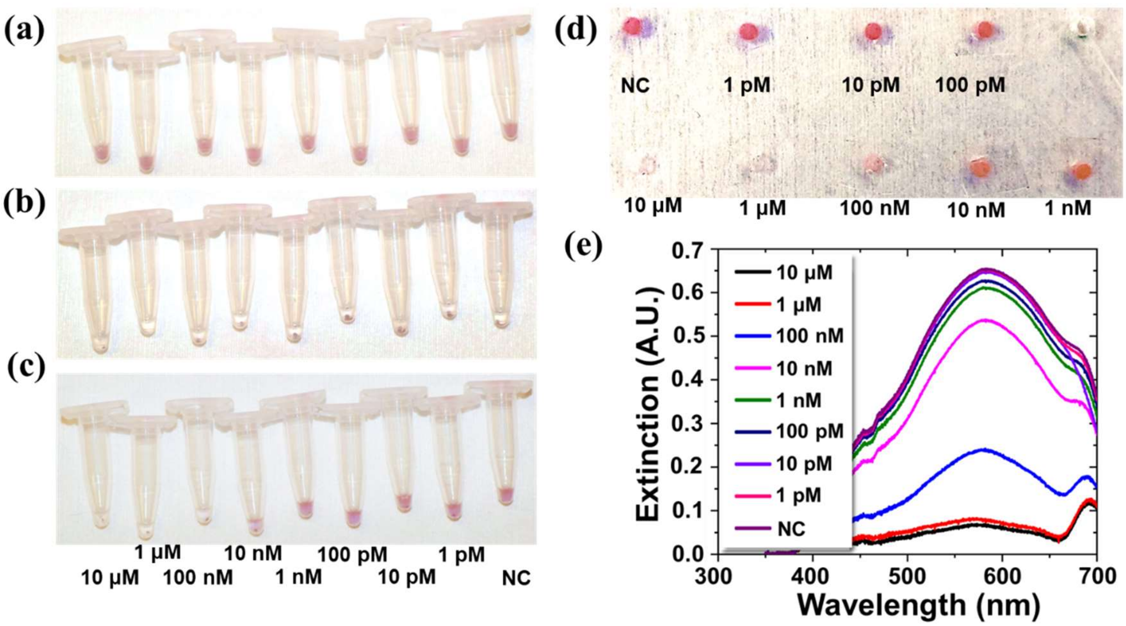


**Figure S13 Images for ICED CBD Detection in 1X PBS with 100 nm AuNPs.** (a-c) Optical images of microcentrifuge tubes (a) after mixing AuNP-CA14 sensor solutions with CBD, molecules (Highlighted step in Figure S5), (b) immediately after mixing AuNP-CA14-CBD with DN21 and centrifugation, and (c) after incubation (20 min) and vortex-mixing. (d) The upper-level liquid from (c) loaded into a PDMS well plate. (e) The spectrometric readout from the well plate.

3. **Study of the Impact of Nanobinders and Sensor Purification on CBD Sensing Performance**

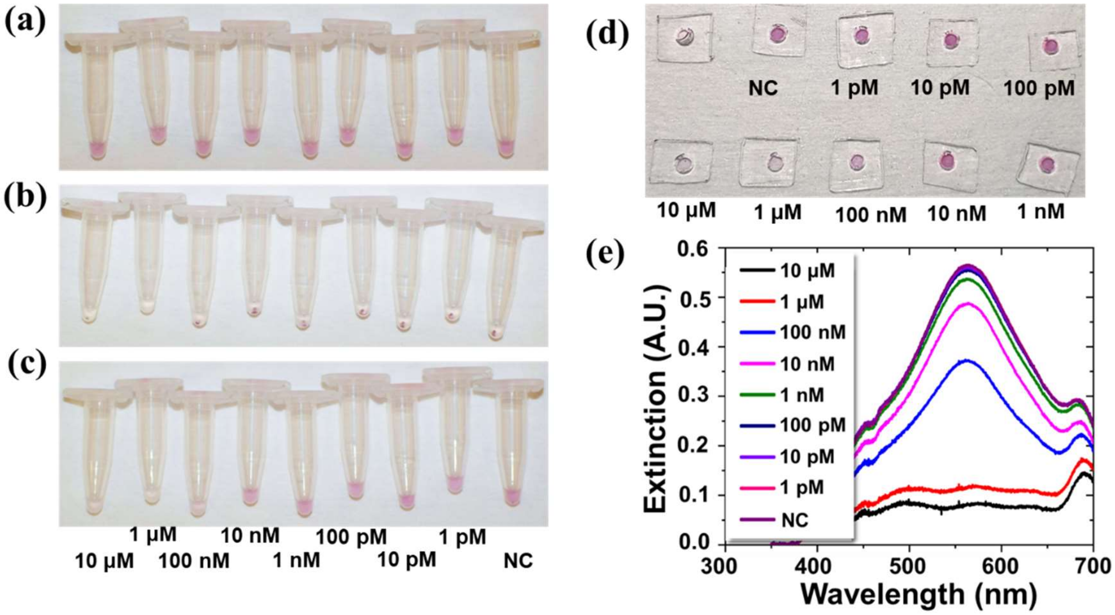


**Figure S14 Images for ICED CBD Detection in 1X PBS with DN18 and 80 nm AuNPs.** (a-c) Optical images of microcentrifuge tubes (a) after mixing AuNP-CA14 sensor solutions with CBD, molecules (Highlighted step in Figure S5), (b) immediately after mixing AuNP-CA14-CBD with **DN18** and centrifugation, and (c) after incubation (20 min) and vortex-mixing. (d) The upper-level liquid from (c) loaded into a PDMS well plate. (e) The spectrometric readout from the well plate.

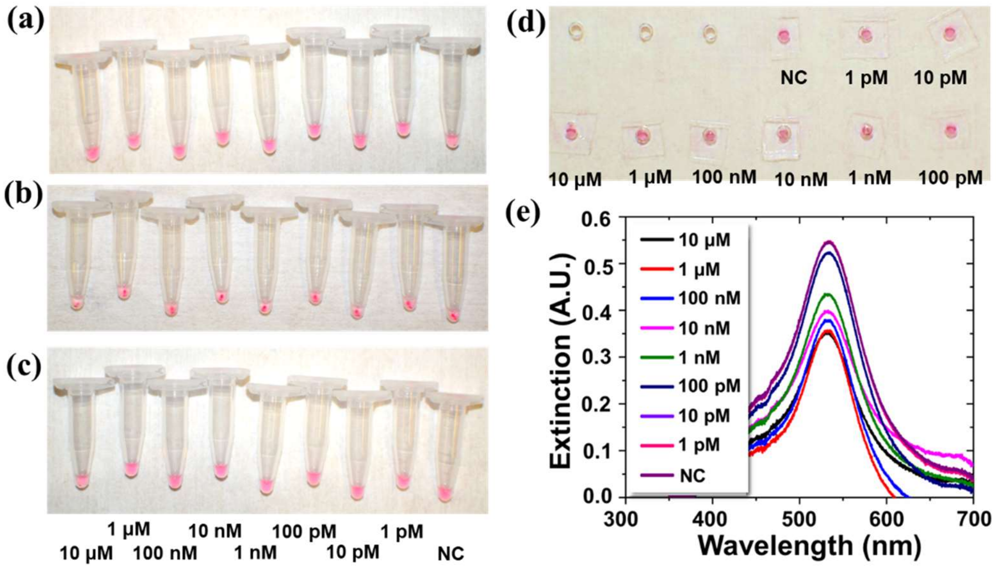


**Figure S15 Images for Unfiltered ICED CBD Detection in 1X PBS with 40 nm AuNPs.** (a-c) Optical images of microcentrifuge tubes (a) after mixing AuNP-CA14 sensor solutions with CBD, molecules (Highlighted step in Figure S5), (b) immediately after mixing AuNP-CA14-CBD with DN21 and centrifugation, and (c) after incubation (20 min) and vortex-mixing. (d) The upper-level liquid from (c) loaded into a PDMS well plate. (e) The spectrometric readout from the well plate.

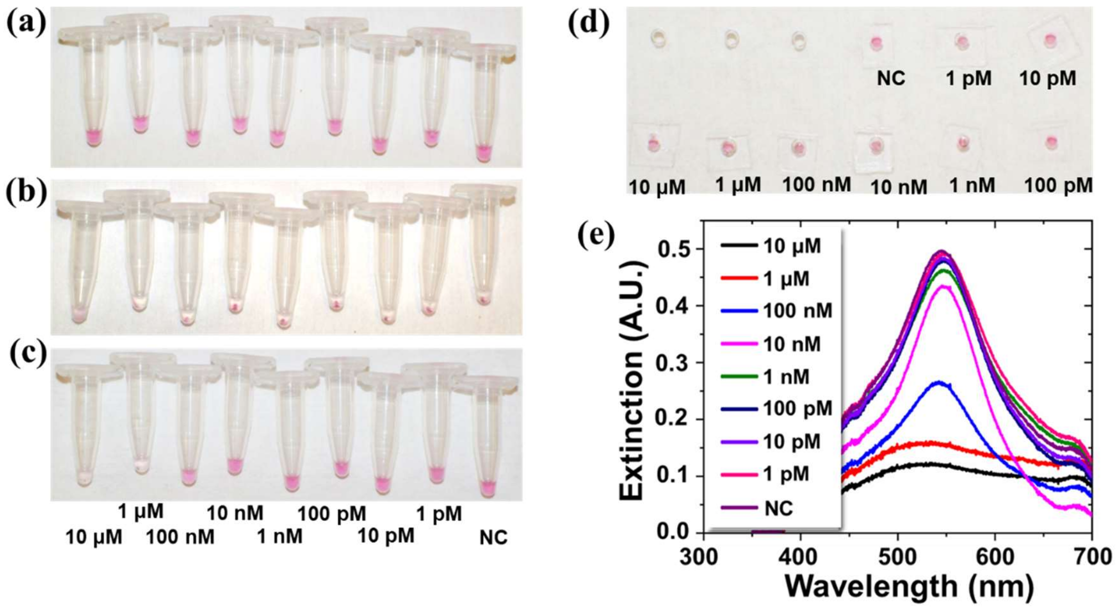


**Figure S16 Images for Unfiltered ICED CBD Detection in 1X PBS with 60 nm AuNPs.** (a-c) Optical images of microcentrifuge tubes (a) after mixing AuNP-CA14 sensor solutions with CBD, molecules (Highlighted step in Figure S5), (b) immediately after mixing AuNP-CA14-CBD with DN21 and centrifugation, and (c) after incubation (20 min) and vortex-mixing. (d) The upper-level liquid from (c) loaded into a PDMS well plate. (e) The spectrometric readout from the well plate.

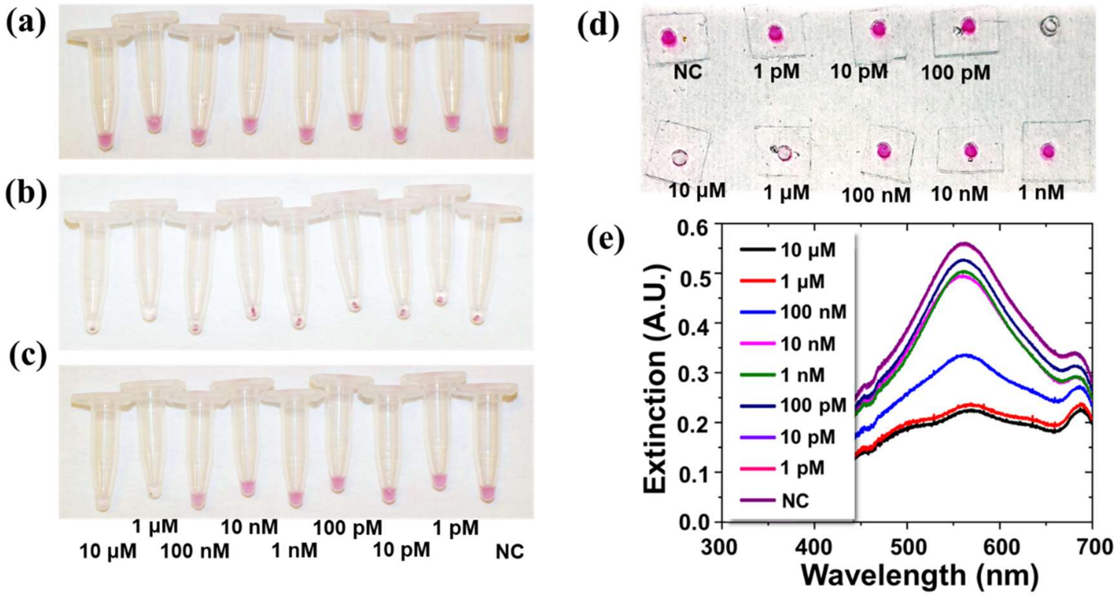


**Figure S17 Images for Unfiltered ICED CBD Detection in 1X PBS with 80 nm AuNPs.** (a-c) Optical images of microcentrifuge tubes (a) after mixing AuNP-CA14 sensor solutions with CBD, molecules (Highlighted step in Figure S5), (b) immediately after mixing AuNP-CA14-CBD with DN21 and centrifugation, and (c) after incubation (20 min) and vortex-mixing. (d) The upper-level liquid from (c) loaded into a PDMS well plate. (e) The spectrometric readout from the well plate.

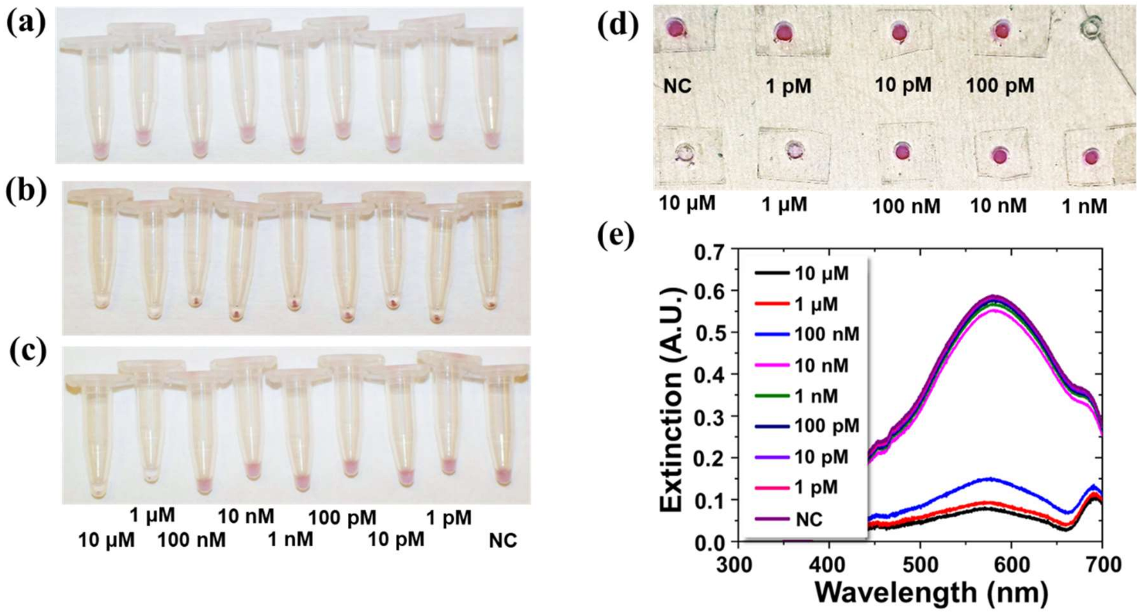


**Figure S18 Images for Unfiltered ICED CBD Detection in 1X PBS with 100 nm AuNPs.** (a-c) Optical images of microcentrifuge tubes (a) after mixing AuNP-CA14 sensor solutions with CBD, molecules (Highlighted step in Figure S5), (b) immediately after mixing AuNP-CA14-CBD with DN21 and centrifugation, and (c) after incubation (20 min) and vortex-mixing. (d) The upper-level liquid from (c) loaded into a PDMS well plate. (e) The spectrometric readout from the well plate.

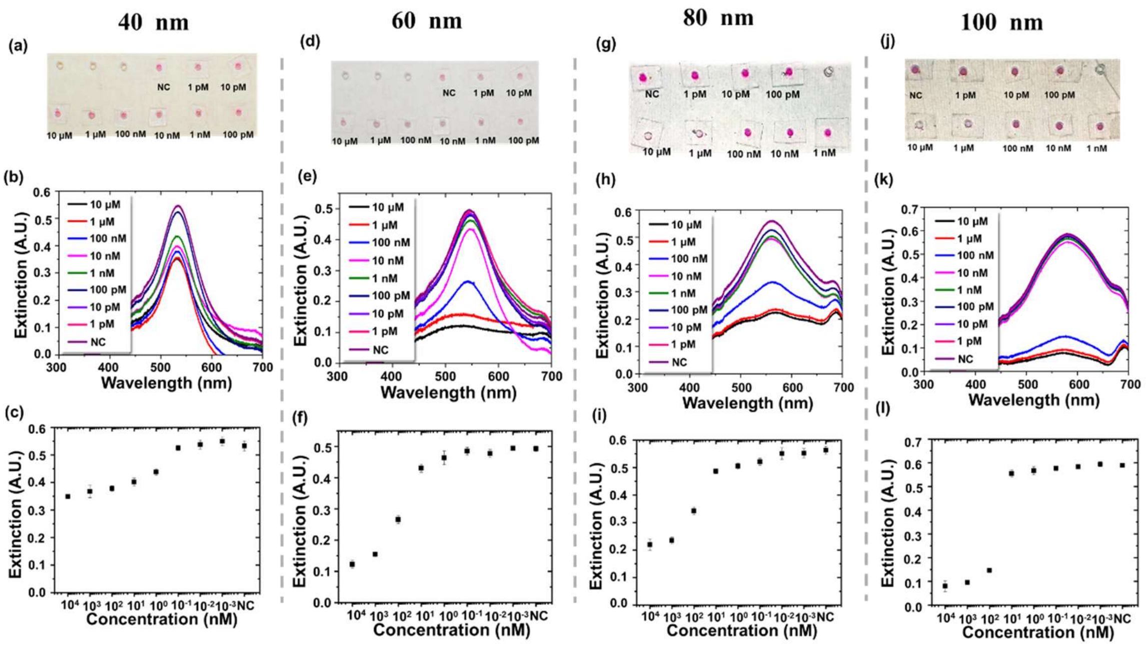


**Figure S19 Summary of the Impact of Nanoparticle Size on CBD Sensing using Unfiltered ICED Scheme in 1 × PBS Buffer.** (a,d,g,j) Optical images of 40, 60, 80, and 100 nm AuNP assay, with the upper-level liquid samples extracted from the microcentrifuge tubes and loaded on a PDMS well plate for CBD detection. (b,e,h,k) Extinction spectra of samples shown in (a)-(d) panel, measured by a lab-based UV-visible spectrometer system. (c,f,i,l) Extinction peak values plotted for AuNP assay using different NP sizes in detecting CBD with concentration from 1 pM to 1 μM. Extinction values were derived from the peak extinction values in (b,e,h,k). In comparison, Figure S10 summarizes the results of different AuNPs where the samples were filtered.

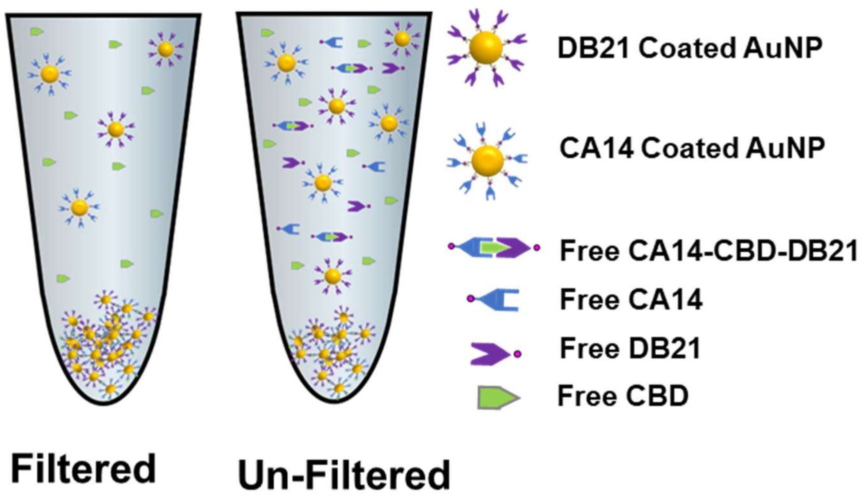


Figure S20 Schematic showing reaction constituents for filtered and unfiltered assay formats.

### 4. CBD and THC Sensing Spiked in Urine and Saliva

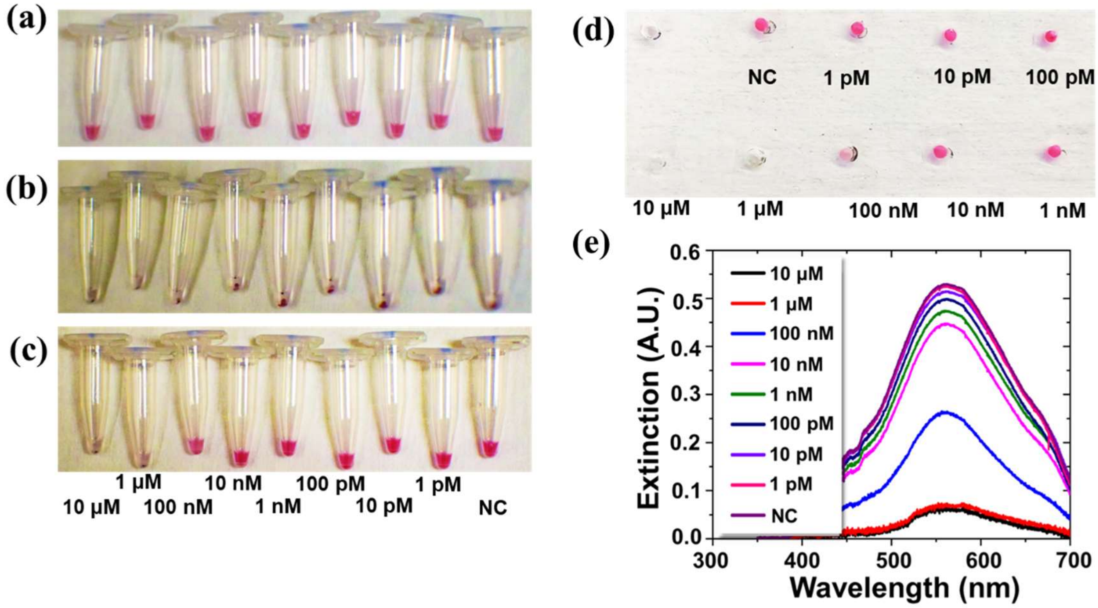


**Figure S21 Experimental results for CBD Detection in 5% Urine Sample using 80 nm AuNPs.** (a-c) Optical images of microcentrifuge tubes (a) after mixing AuNP-CA14 sensor solutions with CBD (in 5% Saliva) , molecules (Highlighted step in Figure S5), (b) immediately after mixing AuNP-CA14-CBD (in 5% Urine) with DN21 and centrifugation, and (c) after incubation (20 min) and vortex agitation. (d) The upper-level liquid from (c) loaded into a PDMS well plate. (e) The spectrometric readout from the well plate.

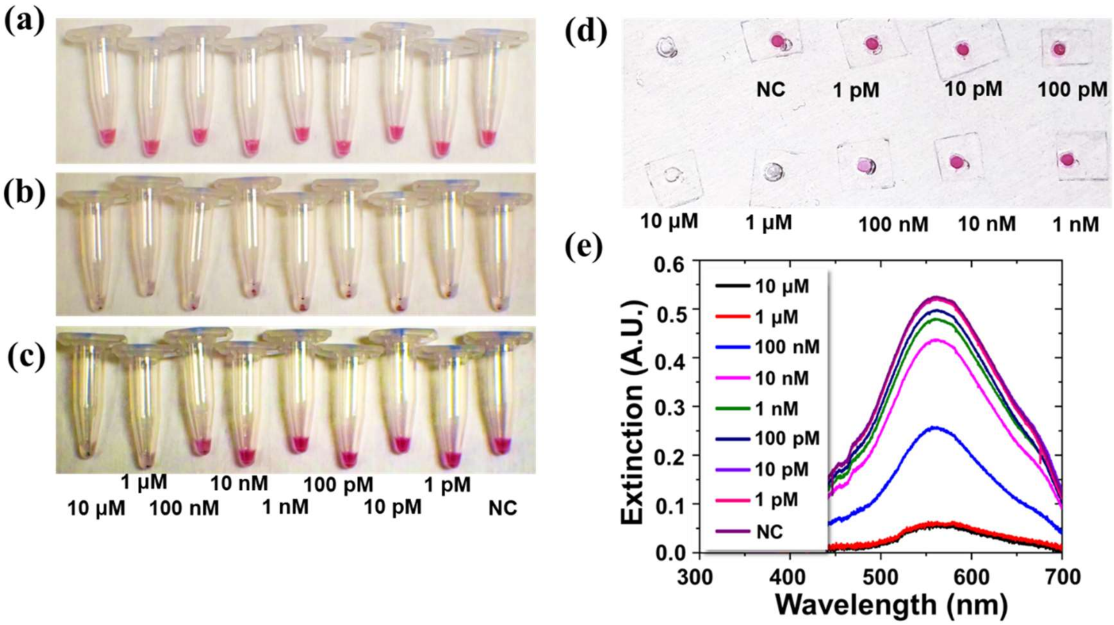


**Figure S22 Experimental results for CBD detection in 5% Saliva Sample with 80 nm AuNPs.** (a-c) Optical images of microcentrifuge tubes (a) after mixing AuNP-CA14 sensor solutions with CBD (in 5% Saliva) , molecules (Highlighted step in Figure S5), (b) immediately after mixing AuNP-CA14-CBD (in 5% Saliva) with DN21 and centrifugation, and (c) after incubation (20 min) and vortex-mixing. (d) The upper-level liquid from (c) loaded into a PDMS well plate. (e) The spectrometric readout from the well plate.

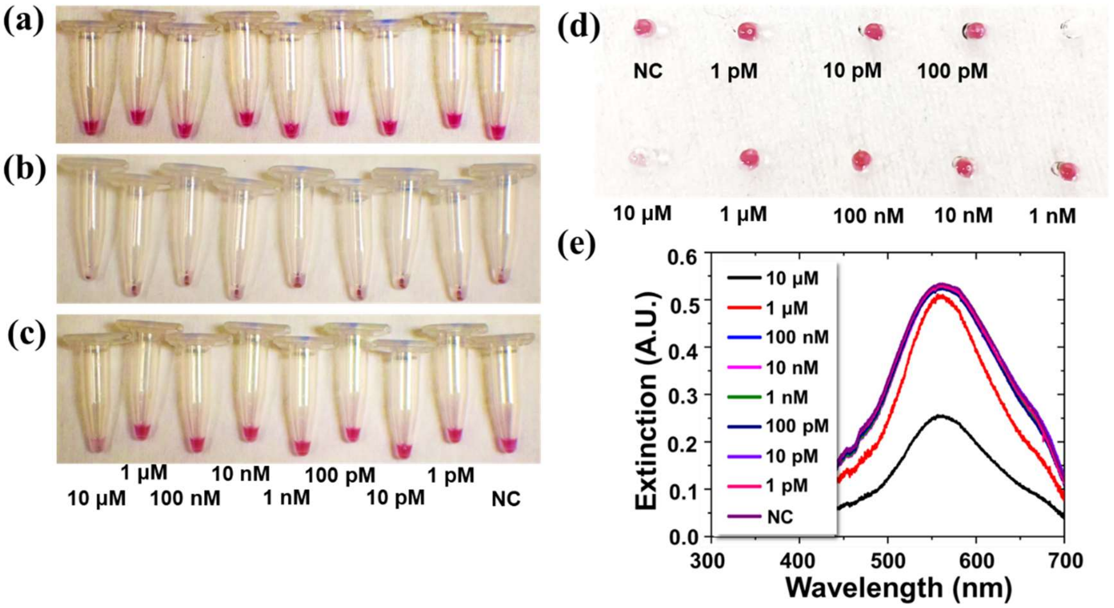


**Figure S23 Experimental results for THC Detection in 5% Urine Sample with 80 nm AuNPs.** (a-c) Optical images of microcentrifuge tubes (a) after mixing AuNP-CA14 sensor solutions with THC (in 5% Urine), molecules (Highlighted step in Figure S5), (b) immediately after mixing AuNP-CA14-THC (in 5% Urine) with DN21 and centrifugation, and (c) after incubation (20 min) and vortex-mixing. (d) The upper-level liquid from (c) loaded into a PDMS well plate. (e) The spectrometric readout from the well plate.

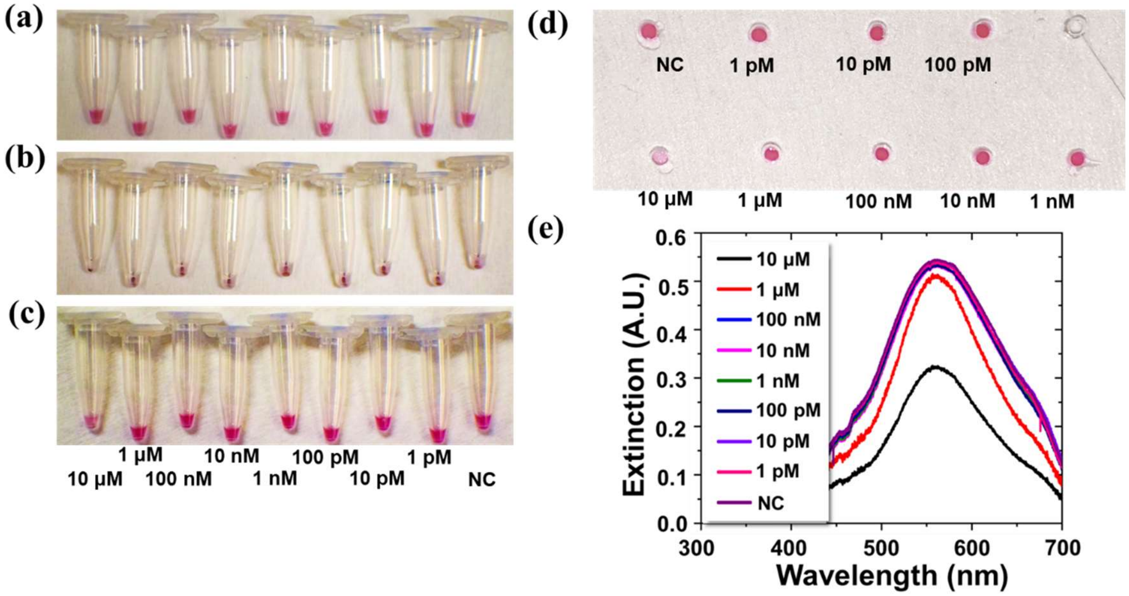


**Figure S24 Experimental results for THC Detection in 5% Saliva Sample with 80 nm AuNPs.** (a-c) Optical images of microcentrifuge tubes (a) after mixing AuNP-CA14 sensor solutions with THC (in 5% Saliva) , molecules (Highlighted step in Figure S5), (b) immediately after mixing AuNP-CA14-THC (in 5% Saliva) with DN21 and centrifugation, and (c) after incubation (20 min) and vortex-mixing. (d) The upper-level liquid from (c) loaded into a PDMS well plate. (e) The spectrometric readout from the well plate.

.

| Steps | Incubation Only | Centrifuge without Pre-incubation | ICED + Filtration | ICED Without Filtration |
|---|---|---|---|---|
| Mixing AuNP: Nano-Binder | 1:2560 | 1:2560 | 1:2560 | 1:400 |
| 2 Stage Filtration | Yes | Yes | Yes | No |
| Pre-Incubation | No | No | Yes | Yes |
| Post Centrifugation | No | Yes | Yes | Yes |
| Post-Incubation | NA | 20 min | 20 min | 20 min |
| Turn around time | 7-8 Hours | 4-5 Hours | 5-6 Hours | 3-4 Hours |
| Detection time | 4 Hours | 20 min | 1 Hour 20 min | 1 Hour 20 min |
| AuNP consumption ( 80 nm AuNP @0.13 nM concentration for 10 samples) | 160 µL | 160 µL | 160 µL | 70 µL |
| Nano-Binders consumption ( @2 µM concentration for 10 samples) | 36 µL | 36 µL | 36 µL | 3 µL |

**Table S1: Summary of regent and time consummation for different detection schemes**

| No | Nanoparticle Size | Dimerization Binder | Medium | Filtration | Centrifugation | Analyte | LOD (nM) All | Measurement Type |
|---|---|---|---|---|---|---|---|---|
| 1 | 40 | DB21 | PBS | Y | Y | CBD | 0.41 | Optical |
| 2 | | | | N | | | 0.45 | Optical |
| 3 | 60 | | | Y | | | 1.20 | Optical |
| 4 | | | | N | | | 2.76 | Optical |
| 5 | 100 | | | Y | | | 0.27 | Optical |
| 6 | | | | N | | | 9.87 | Optical |
| 7 | 80 | | | Y | | | 0.15 | Optical |
| 8 | | | | N | | | 0.22 | Optical |
| 9 | | DB18 | | Y | | | 0.75 | Optical |
| 10 | | DB21 | | | N | | 0.79 | Optical |
| 11 | | | 5% Urine | | Y | | 0.16 | Electronic |
| 12 | | | 5% Saliva | | | | 0.2 | Electronic |
| 13 | | | 5% Urine | | | THC | 1221 | Electronic |
| | | | 5% Saliva | | | | 1123 | Electronic |

Table S2**: Summary of sensing performance under different experimental protocols.**
